# Topographer Reveals Dynamic Mechanisms of Cell Fate Decisions from Single-Cell Transcriptomic Data

**DOI:** 10.1101/251207

**Authors:** Jiajun Zhang, Qing Nie, Tianshou Zhou

## Abstract

Cell fate decisions play a pivotal role in development but technologies for dissecting them are limited. We developed a multifunction new method, *Topographer* to construct a ‘quantitative’ Waddington’s landscape of single-cell transcriptomic data. This method is able to identify complex cell-state transition trajectories and to estimate complex cell-type dynamics characterized by fate and transition probabilities. It also infers both marker gene networks and their dynamic changes as well as dynamic characteristics of transcriptional bursting along the cell-state transition trajectories. Applying this method to single-cell RNA-seq data on the differentiation of primary human myoblasts, we not only identified three known cell types but also estimated both their fate probabilities and transition probabilities among them. We found that the percent of genes expressed in a bursty manner is significantly higher at (or near) the branch point (∼97%) than before or after branch (below 80%), and that both gene-gene and cell-cell correlation degrees are apparently lower near the branch point than away from the branching. *Topographer* allows revealing of cell fate mechanisms in a coherent way at three scales: cell lineage (macroscopic), gene network (mesoscopic) and gene expression (microscopic).

---

Multi-cell organisms start as a single cell that matures through complex processes involving multiple cell fate decision points, leading to functionally different cell types, many of which have yet to be defined^1^. While cellular processes such as proliferation, differentiation and reprogramming are governed by complex gene regulatory programs, each cell makes its own fate decisions by integrating a wide array of signals and executing a complex choreography of gene regulatory changes^2^,^3^. Since the structure of a multi-cell tissue is tightly linked with its function^4^, elucidating the integrative (from gene to cell) mechanism of cell fate decisions is crucial yet challenging.

Single-cell measurement technologies^5^,^6^, which can simultaneously measure the expressions of many genes in a large number of single cells, provide an unprecedented opportunity to elucidate developmental pathways and dissect cell fate decisions. Several algorithms (see a recent review^7^) have been developed to organize single cells in pseudo-temporal order based on transcriptomic divergence and cell states classification. It has been a major challenge to illuminate the dynamic mechanisms of cellular programs governing fate transitions from single-cell data that lacks temporal information^1^. The current methods have mainly focused on identifying trajectories between the most phenotypically distant cell states, and they are usually less robust in reconstructing trajectories from early states towards intermediate or transitory cell states (e.g., Wishbone^8^, Diffusion Pseudotime^9^, Cycler^10^, and CellRouter^11^). Some of the methods have focusing on gaining insights into the regulatory mechanisms driving cell differentiation (e.g., Monocle^12^, ERA^13^, Waterfall^14^, and PIDC^15^), and they seem not to consider how discontinuous, stochastic fate transition events are driven by the dynamic nature of the developmental landscape (which changes in response to activity of gene regulatory networks and extracellular signals) and reflected in the observed increased transcriptional heterogeneity at transition points. In all the existing methods, cell-type dynamics are mainly characterized qualitatively, providing little or none quantitative information on in-depth characterization of complex cellular ecosystems involving cell fate decisions. For a system of multiple cell fate decision points, it has been difficult for the current methods to estimate cell types and their transitions. How fate transitions in the single cell data are related to cell-state gene regulatory networks and the characteristics of transcriptional bursting remain largely unknown.

To overcome the above challenges and to address the important issues on cell fate decisions, we developed *Topographer*, an integrated pipeline that constructs a ‘quantitative’ (e.g., each cell is endowed with ordering and potential) developmental landscape, reveals stochastic dynamics of cell types by estimating both their fate probabilities and transition probabilities among them, and infers dynamic characteristics of transcriptional bursting kinetics along the identified developmental trajectory. In addition, *Topographer* can also both identify various branched (e.g., bi- and tri-) cell-state transition trajectories with multiple branching points from single-cell data and infer networks of marker genes and their pseudo-temporal changes. Together, *Topographer* enables construction of complex cell lineages, resolving intermediate developmental stages, and revealing multilayer mechanisms of cell fate decisions in a coherent way at three different levels: cell lineage, gene network and transcriptional burst (referring to **Supplementary Fig. 1**).

We demonstrated effectiveness of *Topographer* by analyzing single-cell RNA-seq data on the differentiation of primary human myoblasts^12^ while showing applications to other examples in **Supplementary Information**. We identified three known cell types: proliferating cells, differentiating myoblasts and interstitial mesenchymal cells, and constructed a quantitative Waddington’s developmental landscape. By estimating the fate probabilities of these cell types and transition probabilities among them, we found that the probability of transition from the proliferating cell type to the interstitial mesenchymal cell type was approximately twice that of transition from the former to the differentiating myoblast type, and that the fate probability of the differentiating myoblast type was approximately equal to that of the interstitial mesenchymal cell type. We also found that the relative number of the genes expressed in a bursty manner was apparently higher at (or near) the branch point (∼97%) than before or after branch (below 80%). In addition, the mean burst size (MBS) / mean burst frequency (MBF) monotonically decreased / increased before branch but monotonically increased / decreased after branch, with the identified trajectories.

## RESULTs

In order to infer the stochastic dynamics of cell fate decisions from single-cell transcriptomic data, *Topographer* makes the following assumption about the data: the information on the entire development process is adequate, or a snapshot of primary tissue represents a complete developmental process. The overall *Topographer*, a multifaceted single-cell analysis platform, comprises five functional modules: (1) the backbone module (**Fig. 1B**); (2) the landscape module (**Fig. 1C**); (3) the dynamics module (**Fig. 1D**); (4) the network module (**Fig. 1E**); and (5) the burst module (**Fig. 1F**). The backbone module is independent of the remaining 4 modules that depends on the former since they make use of information on cell-state transition trajectories identified in the first module. However, each module achieves an independent function.

**Figure 1.**
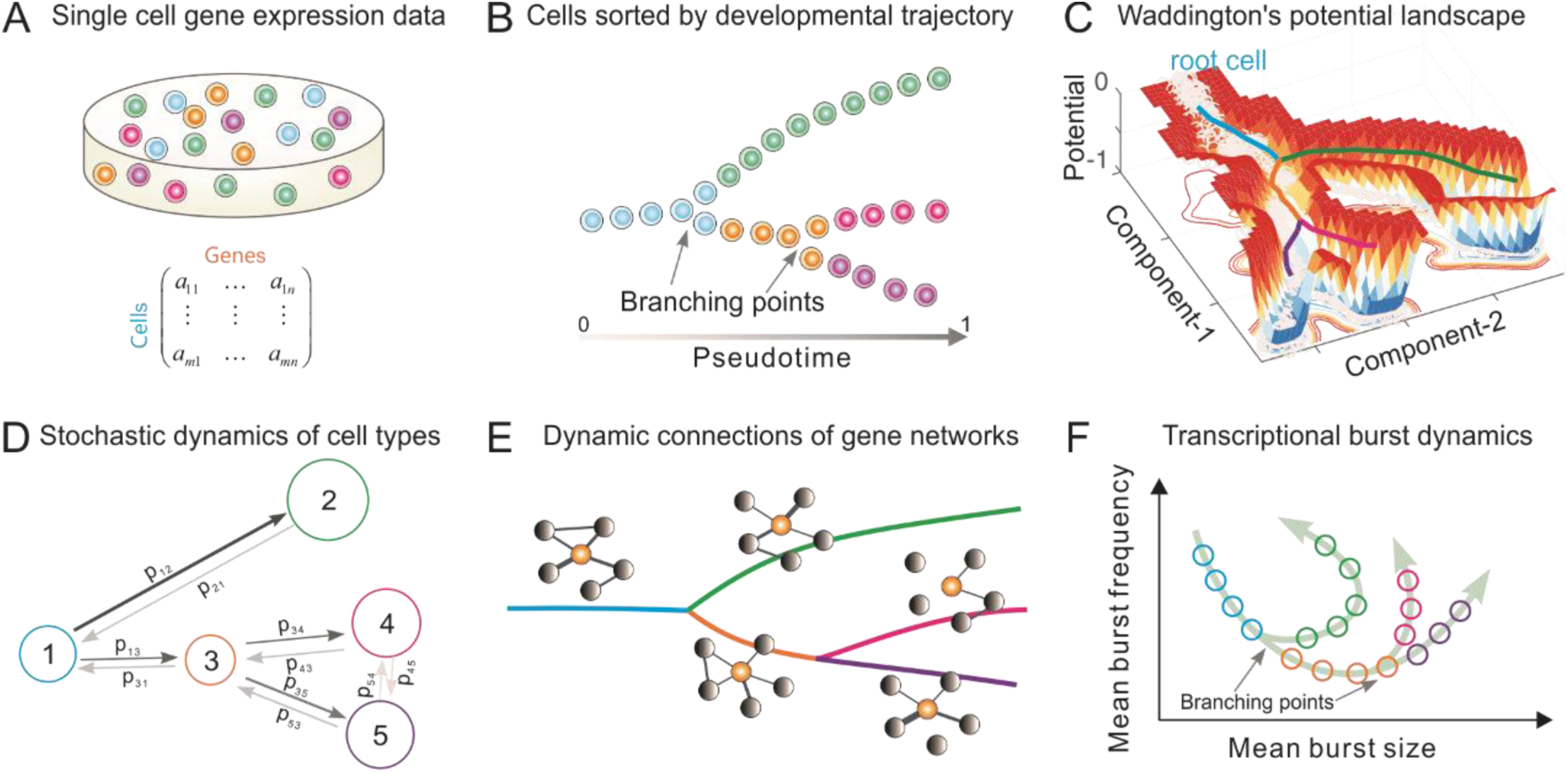
Overview of *Topographer*. *Topographer* comprises five functional modules with each (B, C, D, E, or F) achieving an independent function. (**A**) Single-cell data are represented by a matrix. (**B**) The backbone module identifies the main cell trajectories from the data. (**C**) The landscape module constructs a quantitative Waddington’s developmental landscape, where each cell is endowed with spatiotemporal information (**Online Methods**), and the thick colored lines represent the backbone of cell trajectories identified by the backbone module. This subfigure is not schematic but is plotted using an artificial set of data generated by a toy model (**Supplementary Eq. (24)**). (**D**) The dynamics module reveals stochastic dynamics of cell types by estimating the fate probabilities of cell types and the transition probabilities (indicated by symbols) among them (**Online Methods**), where numbers 1∼5 represent cell types, the size of circle represents that of fate probability, and the thickness of line with arrow represents the size of transition probability. (**E**) The network module infers marker gene networks and their changes along the identified cell trajectories (or along the pseudotime), where the orange ball represents a marker gene, and the thickness of connection line represents the strength of correlation. (**F**) The burst module infers dynamic characteristics of transcriptional bursting kinetics (characterized by both burst size and burst frequency) along the pseudotime, where arrows represent the pseudotime direction.

Two important notes on this method are (1) *Topographer* is unsupervised and needs no prior knowledge of specific genes that distinguish cell fates, and is thus suitable for studying a wide array of dynamic processes involving fate transitions. (2) Except for the backbone module, the other four functional modules of *Topographer* only use the pseudotime information derived in that module (**Online Methods**), so they can also use the result on pseudo-temporal ordering of single cells obtained by other existing algorithms^7^ to achieve their respective purposes. However, the backbone module is established based on a different approach (the following content for details), and has many advantages in contrast to the existing algorithms, e.g., it can identify not only cell-state transition trajectories with multiple branching points but also intermediate or transitory cell states.

Below we introduce each of the five functional modules separately (**Online Methods** gives more details and **Supplementary Information** provides a complete description).

### 1. Identifying the backbone of cell trajectories from single-cell data

The backbone module is a fast and local pseudo-potential-based algorithm. The pseudo-potential here is defined as the negative of the logarithm of a local density function (**Eq. (1), Online Methods**), which aims to identify the backbone of cell-state transition trajectories cross development and find valley floors in a developmental landscape from single-cell data.

Starting from an initial cell (**Fig. 2A**) selected either based on the global minimal pseudo-potential or the prior knowledge, *Topographer* calculates an adaptive step (**Supplementary Eq. (5)**) and searches for pseudo-potential wells on a super-ring (i.e., a high-dimensional circular tube) centered at this initial cell and with the radius equal to the step length (also **Fig. 2A**). In this search method, which clusters cells on super-rings, cluster centers are characterized by a lower pseudo-potential than their neighbors and by a relatively larger distance from points with lower pseudo-potentials (e.g., the only two pseudo-potential wells with ‘green ball’ in **Fig. 2D** are desired), providing the basis of a procedure to find pseudo-potential wells on a super-ring. In this procedure, the number of pseudo-potential wells arises naturally, outliers are automatically spotted, and pseudo-potential wells are recognized regardless of their shape and the dimensionality of the space in which they are embedded. We stress that although there is an analogy between our method and a density-based approach developed originally by Rodriguez and colleagues^16^, the difference is that the former is carried out on a super-ring rather than in the full cell state space. Clearly, if the number of the found pseudo-potential wells (but not including the one found on the ‘reverse’ search direction) is more than one, then this implies the occurrence of branch. The segments linking the center and the newly found pseudo-potential well/or wells on the super-ring can be viewed as part/or parts of the entire developmental trajectory. Similar processes are repeated recursively on sequential super-rings along search directions until no new pseudo-potential wells are found (**Fig. 2B**). By linking all the centers and all the pseudo-potential wells found on super-rings, *Topographer* thus builds a tree-like developmental backbone (**Fig. 2C**). Note that the identified backbone is actually a projection of the pseudo-potential landscape. By projecting every cell onto this backbone (**Subsection 1.2, Online Methods**) and by selecting a root node in the tree (e.g., based on the prior knowledge), *Topographer* thus orders all the single cells in the dataset, and equips each cell with a pseudotime if this root node is set as an initial moment (without loss of generality, the full pseudotime may be set as the interval between 0 and 1).

**Figure 2.**
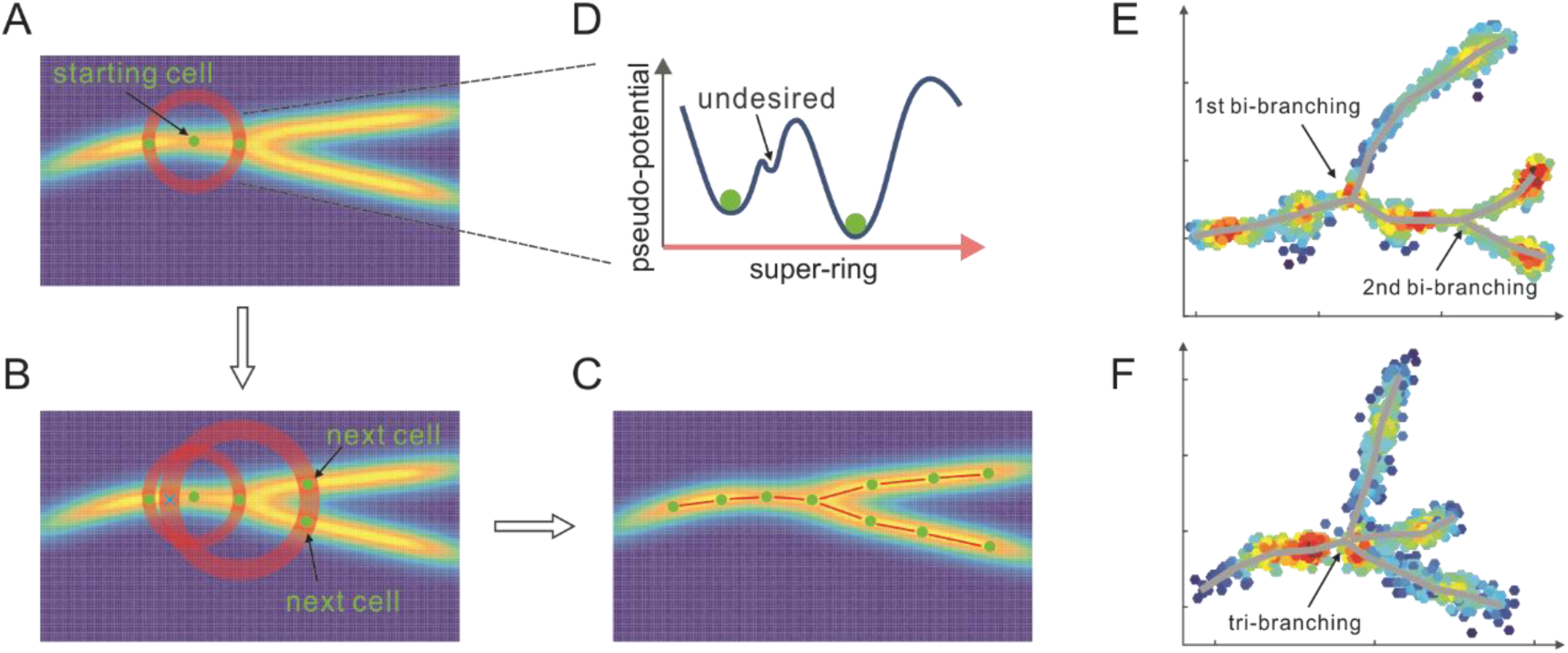
*Topographer* identifies the backbone of branched trajectories from a dataset. (**A, B, C**) A workflow chart (indicated by arrows): *Topographer* first selects an initial cell as the center of a super-ring in the cell state space and searches for pseudo-potential wells on this ring (**A**). Then, *Topographer* repeats recursively on every newly found pseudo-potential well (**B**), where symbol ‘X’ represents a pseudo-potential well found on the reverse search direction, which needs to be excluded in the search process, until no pseudo-potential wells are found, thus building a tree-like backbone of cell trajectories (**C**). Finally, *Topographer* projects every cell onto the backbone, thus ordering all the cells in the dataset. (**D**) shows a super-ring example, where one undesired pseudo-potential well is indicated. (**E**) Bi-branching trajectories identified from an artificial set of data. (**F**) Tri-branching trajectories identified from another artificial set of data.

**Fig. 2E** showed a doubly bi-branched trajectory identified from one simulated dataset, and **Fig. 2F** showed a tri-branched trajectory identified from another artificial set of data. **Fig. 3A** below demonstrated a two-dimensional projection of the de novo cell trajectories identified from single-cell RNA-seq data on the differentiation of primary human myoblasts and **Fig. 3B** demonstrated the evolutions of five marker genes (MYOG, MYF5, MYH2, CDK1 and MEF2C) with branches along the along the pseudotime. **Supplementary Fig. 12** and **Fig. 14** demonstrated results of other two examples, which further showed the power of *Topographer* in pseudo-temporally ordering single cells in single-cell data.

Because of its ability to find pseudo-potential wells on super-rings, *Topographer* can identify developmental trajectories with non-, bi- and multi-branches (referring to **Fig. 1E, F**) (note: a low resolution of experimentally sampling data may lead to tri-branches). In this sense, *Topographer* outperforms previous algorithms, e.g., Waterfall^14^ or Monocle^12^ reconstructs only linear trajectories, and Wishbone^8^ identifies only a single branch point.

**Figure 3.**
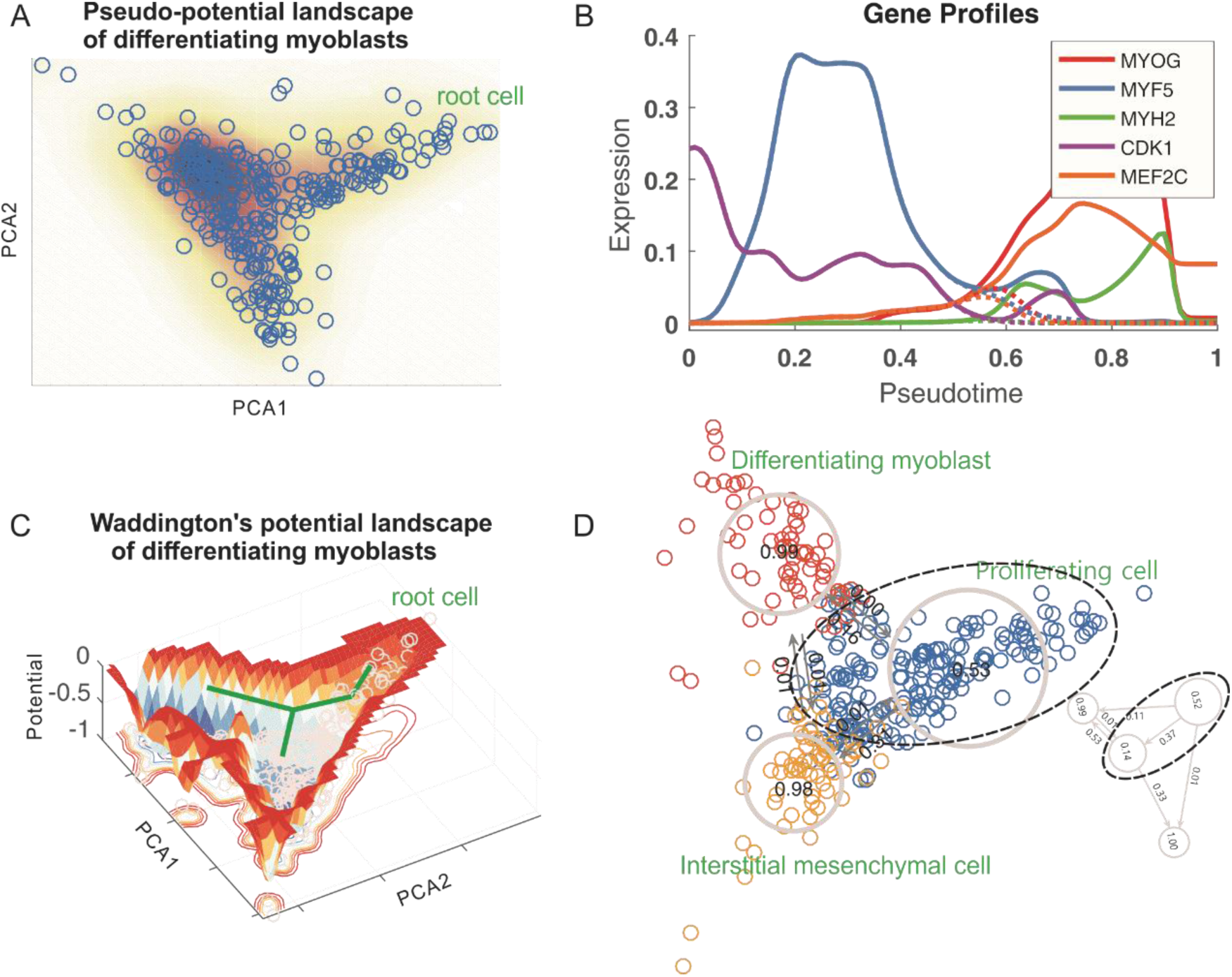
Results obtained by analyzing single-cell RNA-seq data on the differentiation of primary human myoblasts. (**A**) *Topographer* constructs a pseudo-potential landscape, where PCA1 and PCA2 represent components, and every empty circle represents a cell. (**B**) Pseudo-temporal kinetics of five marker genes (indicated by different colors) underlying cell fate decisions, where dashed lines represent the expression levels after branch. (**C**) *Topographer* constructs a Waddington’s potential landscape, where a thick, green line with branch corresponds to the backbone of cell-state transition trajectories identified by the backbone module, and every small, grey circle represents one cell. The normalized potential is shown with the depth of color representing the size of potential. (**D**) *Topographer* reveals stochastic dynamics of cell types along the identified trajectories by estimating both the fate probabilities of cell types (distinguished by colors) and transition probabilities among them. Three known cell types: proliferating cells, differentiating myoblast and interstitial mesenchymal cells, are indicated by dashed ellipses and circles. The large, dashed ellipse shows that the proliferating cell type (top panel) can further be divided into two subtypes (below panel), where the fate probabilities of cell subtypes and the transition probabilities between them are also indicated.

### 2. Constructing a quantitative developmental landscape of single-cell data

The backbone module uses pseudo-potentials to construct the backbone of cell-state transition trajectories, which extracts the information on both branch and cellular ordering from single-cell data, but this kind of potential would not correctly reflect transitions between cells since probability fluxes would exist between them due to cell division, cell death and/or other factors. For example, precursor cells should in principle have higher potentials (**Eq. (8), Online Methods**) in a Waddington’s landscape in contrast to their generations, but if the precursor cells have higher densities, then they have lower pseudo-potentials. Both are apparently inconsistent. In addition, pseudo-potential lacks the temporal information on differentiation or development.

Because of both the above shortcomings of pseudo-potential and the intuition of the Waddington’s potential landscape (in fact, it has extensively been viewed as a powerful metaphor for how differentiated cell types emerge from a single, totipotent cell), the landscape module (an algorithm) is designed to construct a ‘stereometric’ developmental landscape (by ‘stereometric’ we mean that each cell is loaded with spatiotemporal information) in contrast to the ‘planimetric’ backbone identified by the backbone module that uses pseudo-potentials lacking pseudotime. This constructed landscape can provide a more intuitive understanding for the whole developmental process. The principle of the landscape module is simply stated below.

Since single-cell data are noisy due to both cellular heterogeneity and gene expression noise, transitions among the cells scattered randomly in the cell state space can be considered as a random walker (this consideration is inspired by Rosvall and Bergstrom’s work^17^). *Topographer* first constructs a weighted directed graph based on the pseudotime information obtained in the backbone module, and then defines a conditional probability (**Eq. (5), Online Methods**) that the random walker moves from one cell to another with relative weight strengths. Then, *Topographer* estimates the visit probability for each cell by solving a master equation (**Eq. (6), Online Methods**), and determines the potential of every cell in the dataset (**Eq. (8)** with **Eq. (7), Online Methods**). All these potentials are then used to construct a Waddington’s developmental landscape. A dimension reduction^18^ is then used for visualization, the nearest neighbor interpolation is used to fit a landscape function of two variables in a 2-dimension space, and a Gaussian kernel is applied to smooth interpolation (**Subsection 2.2, Online Methods** or **Subsection 3.2.2, Supplementary Information**). In this constructed landscape, each cell is equipped with both potential and pseudotime: two important attributes of a cell. Therefore, the identified backbone of cell-state transition trajectories, which considers pseudo-potentials rather than potentials, can be viewed as an aerial photograph of the constructed Waddington’s potential landscape (comparing **Fig. 3A** with **Fig. 3C)**.

To demonstrate effectiveness of the landscape module, we analyzed two examples: the one for the same set of artificial data used in **Fig. 2E**, with the result demonstrated in **Fig. 1C**, and the other for a set of single-cell data on the differentiation of primary human myoblasts, with the results demonstrated in **Fig. 3C**. It seemed to us that **Fig. 3C** was the first Waddington’s developmental landscape constructed from a realistic set of data (compared with Fig. 5 in Ref. [19], a cartoon). **Supplementary Fig. 13** demonstrated another Waddington’s developmental landscape constructed using single-cell data on the development of somatic stem cells.

**Figure 4.**
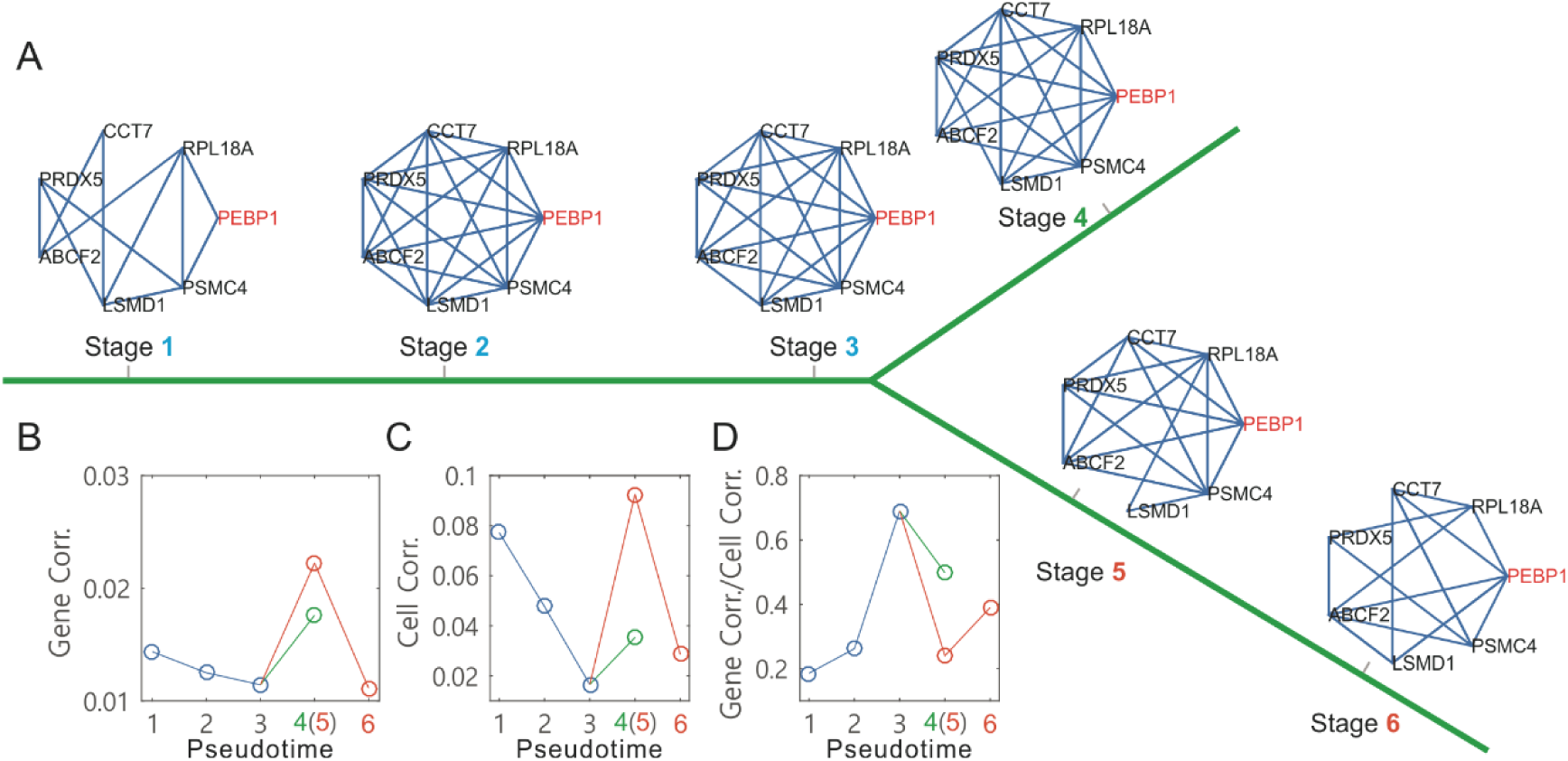
*Topographer* infers dynamic changes in the local connection network of a marker gene along the pseudotime from single-cell transcriptomic data on the differentiation of primary human myoblasts. (**A**) Dynamic changes in a connection network of seven genes along the pseudotime, where the PEBP1 gene (orange) is taken as a core node of neighborhood networks. (**B**) Dynamic changes in the gene-gene correlation degree along the pseudotime before and after branch (different colors), where 6 empty circles correspond to the networks at 6 stages indicated in (A), respectively. (**C**) Dynamic changes in the cell-to-cell correlation degree along the pseudotime before and after branch. (**D**) Dynamic changes in the ratio of the gene-to-gene correlation degree over the cell-to-cell correlation degree along the pseudotime before and after branch.

**Figure 5.**
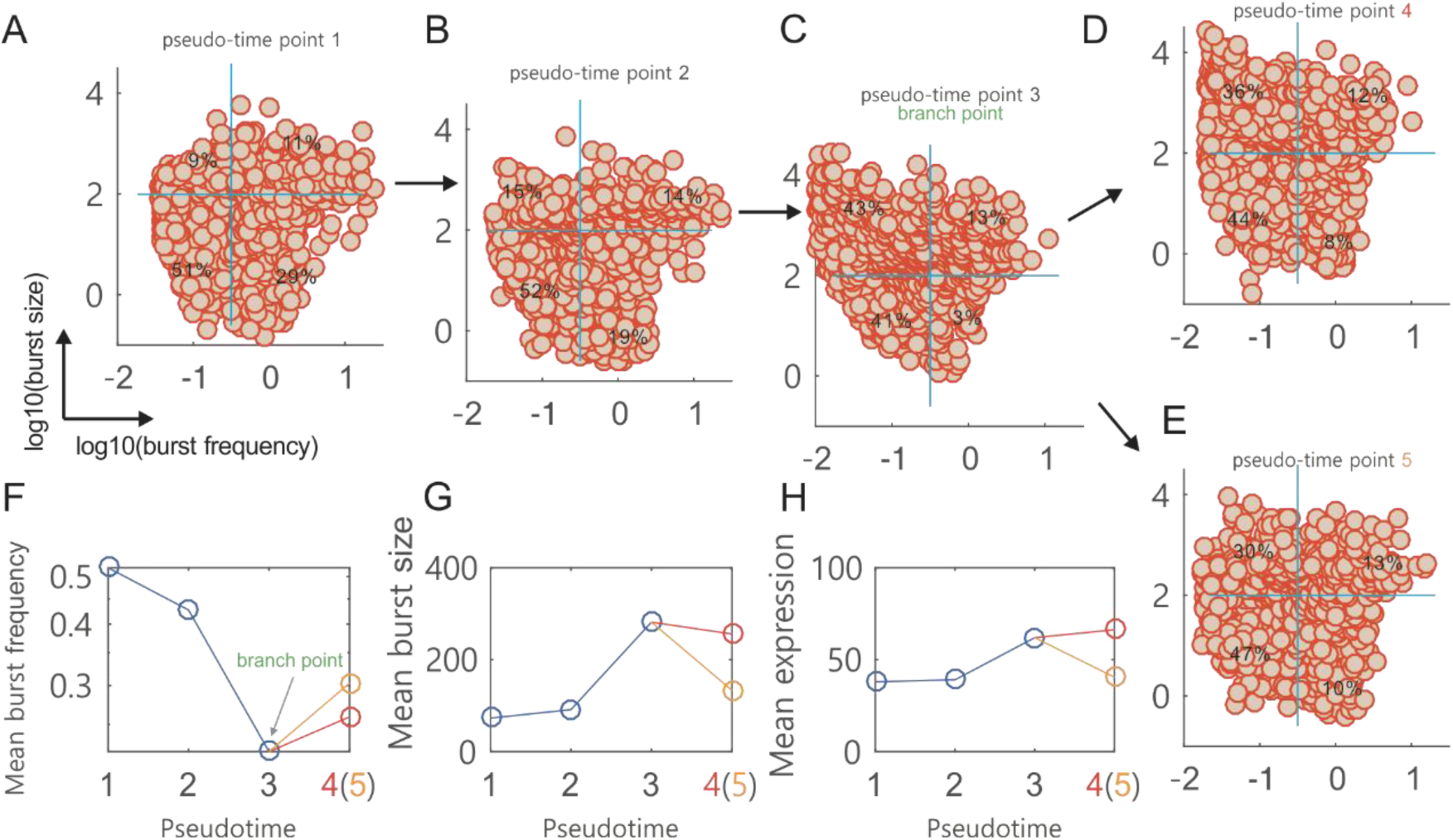
*Topographer* infers dynamic characteristics of transcriptional bursting kinetics along the pseudotime from single-cell RNA-seq data on the differentiation of primary human myoblasts. **(A-E)** Scatter plot of the cells in the logarithmic plane of burst size (BS) and burst frequency (BF) at four pseudotime points, where every circle represents a cell in the dataset. Four percents are indicated in a reference system (two orthogonal blue lines at every subfigure, which correspond to mean BS and BF, respectively). Numbers 4 and 5 actually represent the same pseudotime point. (**F**) Evolution of the mean BF along the pseudotime, where the branching point is indicated and two empty circles after the branching point correspond to (D) and (E), respectively. (**G**) Evolution of the mean BS along the pseudotime. (**H**) Evolution of the mean mRNA expression level along the pseudotime.

It is worth noting that: (1) In contrast to the backbone module that is mainly used to identify a main ‘road’ but ignores “bumpiness” of the road, the landscape module considers both the road (actually a valley floor of the constructed Waddington’s potential landscape) and its bumpiness (reflected by the height of potentials). (2) Both modules can identify cell-state transition trajectories from a dataset, but the former uses pseudo-potentials that rely on neither pseudotime nor cell type whereas the latter uses potentials that depend on both pseudotime and cell type (**Eq. (8)** with **Eq. (4), Online Methods**). (3) Pseudo-potential cannot correctly reflect the motion of a ‘ball’ (i.e., progenitor cell progression) in the constructed Waddington’s potential landscape in which the ball has lower potential at the beginning than at the end, since a lower cell density means a higher pseudo-potential. **Supplementary Fig. 5** shows a difference between potential and pseudo-potential.

### 3. Estimating fate and transition probabilities from single-cell data

Gene regulatory programs underlying cell fate decisions drive one cell type toward another. Quantifying such a transition using single-cell data is challenging due to both cellular heterogeneity and the noise in gene expression in the data.

In order to estimate cell-type dynamics characterized by fate and transition probabilities from single-cell RNA-seq data, it is first needed to determine types of the cells in the dataset. *Topographer* determines cell type according to the following rules: (1) each branch of the identified developmental trajectory is viewed as a cell type with a different branch representing a different cell type; (2) At each branch, the found potential well is taken as a cell subtype with a different potential well representing a different cell subtype. Thus, the number of cell types is equal to the number of branches whereas the total number of cell subtypes is equal to that of potential wells. We will not distinguish cell type and cell subtype unless confusion arises. The cell types determined using this method depend on the shapes of rugged potential wells (prior knowledge can provide additional information in some cases). Therefore, the classification of cell types in this approach is relative rather than “absolute”. For example, in **Fig. 3D**, the proliferating cell type indicated by a dashed ellipse can be further divided into two subtypes. In some situations, a potential well in the constructed Waddington’s potential landscape might not be apparent but still represents a small cell subtype or an intermediate cell state, which may have important biological implications.

In the dynamics module, *Topographer* considers that transitions among the cells scattered randomly in the cell state space is a random walker who randomly moves from a cell state to another, and then estimates two kinds of probabilities: the fate probability for each cell type and the transition probabilities between every two cell types (**Online Methods**). In these estimations, *Topographer* makes use of the cell-state transition trajectories identified in the backbone module.

Specifically, *Topographer* first defines a weight of the directed edge from one cell to another based on the pseudotime (**Eq. (4), Online Methods**), and then uses all the possible weights to estimate the visit probability that the random walker visits a cell in the state space, and further the conditional probability defined as a relative link weight (**Eq. (5), Online Methods**). With these two kinds of probabilities, *Topographer* further estimates the probability that the random walker visits each cell type, and the transition probabilities between every two cell types (**Eq. (9), Online Methods**). These estimations indicate that transitions between cell types are in general not deterministic but stochastic (referring to **Fig. 3D**). In addition, *Topographer* estimates the fate probability of each cell type (**Eq. (12), Online Methods**).

In order to demonstrate stochastic cell-type dynamics estimated by the dynamics module, we again analyzed a simulated data with results shown in **Supplementary Fig. 6**, and a realistic set of single-cell RNA-seq data on the differentiation of primary human myoblasts with results demonstrated in **Figure 3D** (as well as another realistic set of single-cell RNA-seq data on the development of somatic stem cells, with results demonstrated in **Supplementary Fig. 13**). From **Fig. 3D**, we observed that the fate probability (∼0.53) for the proliferating cell type is about the half of that for the differentiating or interstitial mesenchymal cell type (this is not strange since the proliferating cells are root ones) but the fate probabilities for the latter two (∼0.99 and ∼0.98, respectively) are approximately equal. In addition, the proliferating cells differentiate into the differentiating cells at the ∼0.16 probability but the inverse differentiation probability is very small (∼0.001). On the other hand, the proliferating cells differentiate into the interstitial mesenchymal cells at the ∼0.31 probability but the inverse differentiation probability is also very small (∼0.01), implying that the proliferating cells tend to differentiate into the interstitial mesenchymal cells. **Fig. 3D** also showed the fate probabilities of cell subtypes and the transition probabilities between them (low panel).

Apart from the above three main functional modules, *Topographer* can also infer both marker gene networks and their pseudo-temporal changes as well as pseudo-temporal characteristics of transcriptional bursting kinetics. We point out that these inferences can in turn be used to infer whether and when (along the pseudotime) the branches of a developmental trajectory occur.

### 4. Inferring marker gene networks and their pseudo-temporal changes

The network module aims to infer the trend of how marker gene networks dynamically change along the identified cell-state transition trajectories. For this, *Topographer* uses the network neighborhood analysis method^20^ (or **Section 4, Online Methods**) to explore dynamic changes in gene regulatory networks (GRNs) across development.

First, *Topographer* uses GENIE3^21^ to generate a series of GRNs along the pseudotime. Then, based on these GRNs, *Topographer* further analyzes the covariation partners of some particular gene (or genes) using a topological network analysis scheme^22^ that can identify those genes most closely correlated with a given gene (or genes) of interest and most closely correlate to each other (See **Online Methods** for details).

Here, we used the network module to analyze single-cell data on the differentiation of primary human myoblasts, and obtained dynamic changes in the connections of marker gene networks along the identified cell-state transition trajectories (**Fig. 4A**, where the PEBP1 gene is a core node of the networks). From the dependences of mean gene-gene correlation degrees (**Fig. 4B**) and mean cell-cell correlation degrees (**Fig. 4C**) on the pseudotime, we observed that before branch, both degrees were a monotonically decreasing function in pseudotime (the blue line, **Fig. 4B** or **C**), but after branch, each became first monotonically increasing and then monotonically decreasing on one branch (the orange line, **Fig. 4B** or **C**), and monotonically increasing on the other branch (the green line, **Fig. 4B** or **C**). However, the change tendency for the ratio of the gene-gene correlation degree over the cell-cell correlation degree was just opposite to that described above (**Fig. 4D**).

### 5. Inferring pseudo-temporal characteristics of transcriptional bursting kinetics

Transcription occurs often in a bursty manner, and single-cell measurements have provided evidence for transcriptional bursting both in bacteria and in eukaryotic cells^23^. By analyzing a simplified stochastic model of gene expression, previously Xie, e al showed^24^ that the number of mRNAs produced in the bursty fashion following a Gamma distribution determined by two parameters: MBF (i.e., the mean number of mRNA production bursts per cell cycle), and MBS (i.e., the average size of the mRNA bursts).

There is great interest in analyzing single cell data to understand the transcriptional changes that occur as cells differentiate and the genes and regulatory mechanisms controlling differentiation processes and cell-fate transition points^2^,^25^. The burst module is designed to infer the trend of how transcriptional bursting kinetics dynamically changes across development. For this, *Topograph*er uses the maximum likelihood method^26^ to infer the two parameters of MBF and MBS from single-cell RNA-seq data (**Section 5, Online Methods**), thus revealing dynamic characteristics of transcriptional bursting kinetics before branch, near the branching point and after branch of the developmental trajectory.

We used the burst module to analyze single-cell data on the differentiation of primary human myoblasts. **Fig. 5A-E** showed how the cells at four pseudotime points (two before branch, one at branch point and one after branch) were distributed in the logarithmic plane of burst frequency (BF) and burst size (BS). A reference system (two indicated orthogonal blue lines: the horizontal line for BF and the vertical line for BS) was used to guide visual comparison between the rates (i.e., the indicated percents) of cell numbers over the total cell number at a particular pseudotime point. The four quadrants of the reference system clearly showed how the genes in the dataset were expressed, e.g., the fourth quadrant showed that the genes were expressed in a manner of high frequency (i.e., the BF is more than 0.33) and small burst (i.e., the BS number is less than 200). We observed that the genes expressed in a bursty manner (i.e., the other three cases except for the case in the fourth quadrant) were more at the branching point (97%) than before or after branch (approximate or below 80%). In other words, the percent of the genes expressed with high frequency and small burst was apparently lower at the branching point. From these figures, we can conclude that during the differentiation of primary human myoblasts, there are more genes expressed in a bursty manner at the branching point than before or after branch.

From the dependences of MBF and MBS on the pseudotime (**Fig. 5F, G**), we observed that there were apparently different change trends before and after branch. **Fig. 5H** showed the dependence of the mean mRNA expression level on the pseudotime, demonstrating a change tendency opposite to that of MBF. Although fundamentally similar to the change trend of MBS on the whole, the mean mRNA level changed almost stably with the pseudotime for one branched trajectory (referring to the top line after branch in **Fig. 5H**). These three subfigures implied that MBF or MBS can be taken as a better indicator of the branch occurrence than the mean mRNA expression level.

## DISCUSSION

We have developed a computational pipeline – *Topographer* for construction of developmental landscapes, identification of de novo continuous developmental trajectories, and quantification of fate transitions. One unique feature of *Topographer* is its capability of characterizing both transcriptional bursting kinetics and changes in connections of marker gene networks along developmental trajectory. When identifying the backbone of cell-state transition trajectories from single-cell data, *Topographer* was robust to the noise in the dataset (**Supplementary Fig. 8**-**10**). When applied to the differentiation of primary human myoblasts, *Topographer* first constructed an intuitive developmental landscape for an order and timing of events that closely recapitulated previous studies of this system. In addition, it estimated the fate probabilities for cell types and the transition probabilities between them. Together, the results suggested that the fate transition during the differentiation of primary human myoblasts occurred in a probabilistic rather than deterministic manner, and the transitions between cell types might be unidirectional and bidirectional. These two new insights challenge the traditional view that the development of primary human myoblasts was tree-like or that the process from a predecessor to its generations was both deterministic and unidirectional^5^.

When ordering single cells, *Topographer* (like existing methods in the literature) needs to assume sufficient number of cells in the dataset because the backbone module is established essentially based on the estimation of cell density. A small number of cells (e.g., less than 100), would lead to inaccuracy of finding pseudo-potential well /or wells on a super-ring in the backbone module. As more cells can simultaneously be measured^22^, the accuracy of *Topographer* will improve. In principle, *Topographer* can also be used to analyze other single-cell data such as mass cytometry data^27^ and single-cell PCR data^28^.

Cell fate decisions may involve hierarchy of cell types including intermediate cell states or cell subtypes. Identifying such (e.g. rare) sub-cell types is important yet challenging. *Topographer* has shown its ability to identify cell subtypes, which may correspond to shallow or small potential wells in the constructed developmental landscape (right below, **Fig. 3D**). Moreover, *Topographer* can estimate the fate probability of each identified cell subtype and the transition probabilities between every two identified cell subtypes (right below, **Fig. 3D**), which is one main advantage of *Topographer* compared to many existing methods^7^. In particular, *Topographer* enables identification of non-, bi- and multi-branches (**Fig. 2C, D**), one main function that have been under developed in the existing algorithms^7^ including Wishbone^8^.

It is worth noting that *Topographer* provides a general framework connecting three interplayed major components based on single-cell data: cell lineage committing dynamics (macroscopic), gene network dynamics (mesoscopic) and transcriptional bursting kinetics (microscopic). First, *Topographer* provides useful information on their relationships that are implied by the pseudotime, but this kind of time only reflects the impact of the former on each of the latter two. The issues of how and in what degree the inferred gene connection networks or/and transcriptional bursting kinetics influence or/and determine cell fates in the underlying developmental process, remain unexplored. In order to study the relationship between the mesoscope / microscope and the macroscope, a possible way is to establish the so-called balance equation^28^. Second, in order to estimate the fate probabilities of cell types and the transition probabilities between them (**Eq. (12)** and **Eq. (9), Online Methods**, respectively), *Topographer* makes an assumption, that is, it assumes that the transition from one cell to another along a cell-state transition trajectory is linear (**Eq. (4)** in **Online Methods** or **Supplementary Eq. (6)**). In many cases, such transition may be nonlinear since, e.g., cell-cell communication through signal molecules. Third, the traditional Waddington’s landscape (e.g., tumor pathobiology^29^) is often used as an intuitive tool to describe differentiation process through the trajectory of a ball into branching valleys with each representing a developmental state^30^. *Topographer* uses the potential of each cell to quantify developmental landscape, which allows estimation of the transition probabilities between cell types and their fate probabilities to characterize cell lineage committing dynamics. These probabilities have physical meanings as they actually represent the Krammer escape rates^31^ between potential wells. However, how cell fate decisions including cell-state dynamics are related to Krammer escape rates remain unclear. Fourth, based on the transition probabilities between cell types, one can establish a model of cell population dynamics (referring to **Supplementary Eq. (21)**), and further study stochastic state transitions from a dynamical-system perspective..

Finally, using “relatively smaller pseudo-potential and relatively larger distance” (**Online Methods**) as a rule in backbone module is a robust approach in finding cell trajectories (referring to **Supplementary Fig. 8**-**10**); In our method, the transitions among cells are considered as a random walker who moves randomly between the data points scattered in the cell state space. These two ground rules used in *Topographer* can be viewed as new principles of mining single-cell data to uncover mechanisms of cell fate decisions.

## Methods

Methods, including statements of data availability and any associated accession codes and references, are available in the online version of the paper.

*Note: Any Supplementary Information and Source Data files are available in the online version of the paper.*

## Acknowledgments

We thank Mr. Xiaoming Lai’s partial computer program codes for the manuscript. This work was partially supported by National Natural Science Foundation of China (11775314, 91530320), and National Key Basic Research Program of China (2014CB964703).

## AUTHOR CONTRIBUTIONS

J.Z. and T.Z. conceived and designed the work. J.Z. carried out computer implementation and data analysis, J.Z., Q.N. and T.Z. interpreted the simulation results. T.Z. supervised the project, J.Z. and T.Z. wrote the original manuscript, and Q.N. contributed to the writing of final manuscript.

## COMPETING FINANCIALINTERESTS

The authors declare no competing financial interests.

## Online Methods

The overall *Topographer*, a multifunctional algorithm, comprises five functional modules: the backbone module, the landscape module, the dynamics module, the network module, and the burst module. Main details of these modules are separately stated below and the complete description including data pre-processing is given in **Supplementary Information**.

### 1. The backbone module identifies the backbone of cell-state transition trajectories from single-cell data

Assume that there are *m* cells and *n* genes in single-cell RNA-seq data of interest, which can in principle be represented as *m* points in the *n* -dimensional space (*X*) of gene expression (called the cell state space for convenience).

the backbone module aims to identify the backbone of cell-state transition trajectories across development from the dataset. The essence is to find valley floors in a developmental landscape. Specifically, *Topographer* finds valleys with local minimal pseudo-potentials, where pseudo-potential is defined as

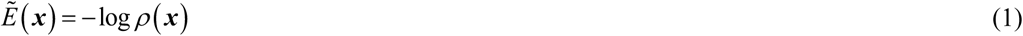

with

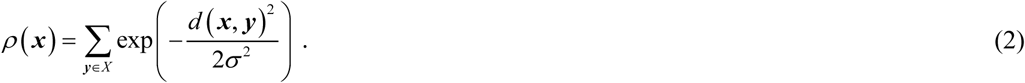

In **Eq. (2)**, *d* is the Euclidean distance between two state points ***x*** and ***y*** in the cell state space *X* (note: other kinds of distances are also suitable for *Topographer*). Note that *ρ* represents the local cell density, mostly accounting for the number of cells in a neighborhood defined by *σ*.

Roughly speaking, *Topographer* starts by cell state *x*_0_ (i.e., an initial cell) and then searches for pseudo-potentials wells on super-rings (which are actually circular tubes in the cell state space) by recursively applying an extended version of the cluster algorithm^16^. Finally, all the centers of the super-rings are represented in a tree, *T*. Main details are stated below and more details are given in **Supplementary Information**.

#### 1.1 Constructing a developmental tree

Starting by an initial cell that has the global minimal pseudo-potential or by the cell that the user chooses according to the prior knowledge, *Topographer* calculates an adaptive radius or an adaptive step length (**Subsection 2.1.4, Supplementary Information**) and searches for potential wells on a super-ring centered at this cell and with the radius (referring to **Fig. 2A**). The search method (called the pseudo-potential well search algorithm) is based on the idea that cluster centers on the super-ring are characterized by a lower pseudo-potential than their neighbors and by a relatively larger distance from points with locally lower pseudo-potentials. Specifically, *Topographer* first defines

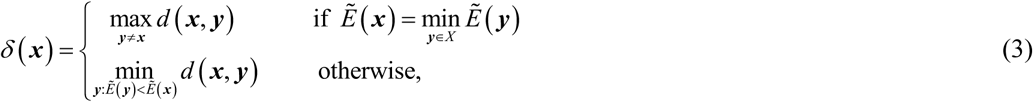

and then finds local pseudo-potential well /or wells on the super-ring, based on the combination of relatively smaller 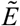 and relatively larger *δ*. Therefore, there is an analogy between the pseudo-potential well search algorithm and a density-based approach developed originally by Rodriguez and colleagues^16^. The segments linking the center and the potential wells found on the super-ring can be taken as approximate part /or parts of the entire developmental trajectory.

Then, taking every found pseudo-potential well as the center of a new super-ring with a new adaptive radius, *Topographer* performs similar calculations as at the previous step, thus finding pseudo-potential well /or wells on this new super-ring. Again, the segments linking the new center and the newly found pseudo-potential wells on the new super-ring can be taken as other approximate part /or parts of the entire developmental trajectory. This process is repeated until no new pseudo-potential wells are found. By linking the cluster centers, *Topographer* thus builds a tree-like developmental backbone, which is actually composed of valleys.

Note that for a super-ring’ center rather than the starting point, the newly found valleys would include valleys on the “reverse direction” in the processes of searching for local pseudo-potential wells on super-rings, which are not expected in our algorithm. To handle such an exception, *Topographer* excludes those valleys that are too close to the found valleys. In addition, any two newly found valleys with the distance of smaller than the step length are merged by discarding the valleys with larger pseudo-potentials. Such a treatment may greatly improve the algorithm’s robustness against the noise in the dataset (referring to **Supplementary Fig. 8**-**10**).

Also note that a complete valley floor is constructed by terminating the recursive process for some super-ring on which no desired pseudo-potential wells can be found. Since no loops are assumed to exist in the developmental trajectory, there is definitely a boundary, implying that the search process necessarily stops within finite steps.

After the above search process is completed, all the found pseudo-potential valley floors are represented in an undirected acyclic graph (a tree with branches). To achieve better accuracy and coverage, *Topographer* refines a pseudo-potential valley tree by searching for pseudo-potential well /or wells on the line linking two centers on every edge of the tree (referring to **Supplementary Fig. 9, 10**). To that end, *Topographer* finishes construction of the backbone of a developmental tree from a given set of single-cell data.

#### 1.2 Cell projection and pseudotime assignment

After constructing a developmental tree, *Topographer* then projects every cell point in the cell state space onto some edge of the tree according to the shortest distance principle (i.e., the perpendicular distance from the cell point to the edge is shortest). Thus, every cell has its unique relative position in the identified backbone (or in the constructed tree).

Next, *Topographer* assigns a pseudotime for every cell in the dataset. Before that, however, it is needed to determine a root node in the constructed tree. *Topographer* chooses a root cell in such a manner that the distances between this cell and those cells that are initially set according to, e.g., the prior knowledge, are as short as possible. An initial pseudotime is first assigned to this root node. Every other cell in the dataset is then assigned in order with a pseudotime according to its relative position in the constructed tree. Without loss of generality, the full pseudotime may be set as the interval between 0 and 1.

### 2. The landscape module constructs a quantitative Waddington’s developmental landscape of single-cell data

#### 2.1 Calculation of cell potential

After the backbone of a developmental trajectory has been identified and every cell has been endowed with a pseudotime value, the landscape module first estimates the potential of every cell in the dataset and then uses these potentials to construct a quantitative developmental landscape. It is expected that the potential to be introduced can be avoid shortcomings of the pseudo-potential as pointed out in the main text. For this estimation, *Topographer* analogizes transitions between cells at distinct stages of the differentiation process to a random walker who moves randomly between the data points that are randomly scattered in the cell state space. This analogy, which is inspired by Rosvall and Bergstrom’s work^17^, is reasonable due to both cellular heterogeneity and gene expression noise in the dataset. In addition, it is important that *Topographer* uses the pseudotime information to construct a weighted directed graph **W**.

Specifically, *Topographer* defines the weight of the directed edge from cell *α* to cell *β* as

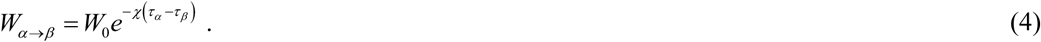

(**Supplementary Information** gives a reason for this definition), where *τ*_*α*_ and *τ* _*β*_ represents the pseudotime points for cells *α* and *β* respectively, and positive constant *χ* represents a linearly changing rate that cell *α* transitions to cell *β* (this setting implies an assumption, i.e., the evolutional process from one cell to another along the pseudotime is assumed to be linear). The setting of the *χ* value in general depends on the dataset under consideration (**Subsection 3.2.1** in **Supplementary Information** gives a simple discussion) but it may be set as 30 in our cases. It is worth pointing out that the weight defined in such a manner has used the information on the pseudo-temporally ordered cell trajectories, which is a key for the entire calculation.

Then, in order to estimate cell visit probability on a random walk, *Topographer* defines a conditional probability that the random walker moves from cell *β* to cell *α* as the relative link weight, given by

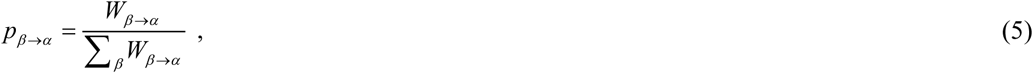

which is apparently independent of initial *W*_0_. If the stationary visit probability of cell *α* is denoted by *p*_*α*_, then *p*_*α*_ can in principle be derived from a recursive system of the form

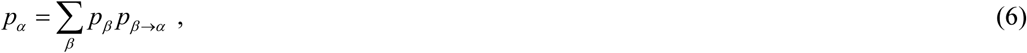

which represents the probability that the random walker visits the *α* cell from all the other possible cells. Note that **Eq. (6)** is actually a master equation^31^ and can efficiently be solved with the power-iteration method^32^. However, to ensure that the unique solution of this equation is independent of the starting node in the directed network, the random walker instead teleports to a random node at a small rate *ε* with 0 < *ε* < 1. In addition, to obtain more robust results that depend less on the teleportation parameter *ε*, it is most often to use teleportation to a node proportional to the total weight of the links to the node^17^. Because of these considerations, the resulting stationary visit probability for cell *α* is modified as

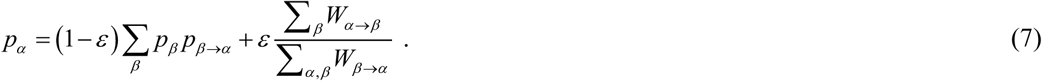

Finally, *Topographer* quantifies the potential of every cell in the dataset, according to

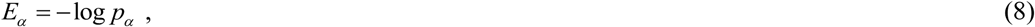

where *p*_*α*_ is given by **Eq. (7)**. Apparently, the potential defined in such a manner has again made use of the information on the identified cell-state transition trajectories due to **Eq. (4)**. We point out that the potential of a cell depends on pseudotime but the pseudo-potential lacks the information on pseudotime.

#### 2.2 Scatter plot of developmental landscape

After all the cells in the dataset have been equipped with potentials, all these potentials are then used to construct a Waddington’s potential landscape for the developmental process. The method is stated as follows. First, dimension reduction is needed for visualization (the tSNE method^18^ or the PCA method^33^ may be used to achieve this purpose). In general, dimension reduction cannot explicitly reflect the information on coordinates in a visualized landscape, e.g., PCA1 and PCA2 in **Fig. 3C** do not actually represent components in the dataset. Second, *Topographer* uses the nearest neighbor interpolation method to perform interpolation on a 3-dimensional scattered data set. Specifically, *Topographer* uses ScatteredInterpolant (a function of the MATLAB software) to establish the corresponding relationships between a set of points, (*x, y)*, and a set of cell potentials, *E*. These relationships, denoted by *E = F (x, y)*, in principle define a curved surface in the 3-dimensional space for the developmental landscape, which in return passes through all the sampling points in the space under consideration. *Topographer* then uses the nearest neighbor interpolation to evaluate this surface at any query point (*x*_*q*_, *y*_*q*_), thus obtaining an interpolating value of every known potential given by **Eq. (8)**, i.e., *E*_*q*_ = *F* (*x*_*q*_, *y*_*q*_). Third, a Gaussian kernel is used to smooth interpolation. Finally, the identified developmental trajectory is drawn on the constructed developmental landscape (referring to the thick colored line in **Fig. 1A** or the thick green line in **Fig. 3C**).

We point out that pseudo-potential cannot correctly reflect the motion of a ‘ball’ in the constructed Waddington’s potential landscape in which the moving ball has lower potential at the beginning than at the end, since a lower cell density implies a higher pseudo-potential according to definitions. **Supplementary Fig. 5** shows a difference between potential and pseudo-potential.

### 3. The dynamics module estimates fate probabilities of cell types and transition probabilities between them from single-cell data

#### 3.1 Determining cell types

Cell-type dynamics can be characterized by fate and transition probabilities. In order to estimate these probabilities, it is first needed to determine the types of cells in the dataset. For this, *Topographer* adopts the following rules: First, each branch in the identified developmental trajectory is defined as one cell type, and a different branch is defined as a different cell type. Then, each potential well on each branch is defined as one cell subtype, and a different potential well is defined as a different cell subtype. These definitions imply that the number of cell types is equal to that of branches whereas the number of cell subtypes is equal to that of potential wells. It should be pointed out that the cell type determined in such a manner is not unique but depends on the choice of 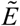 and *δ* (their respective definitions above). In the following, we will not distinguish cell type and cell subtype unless confusion arises.

#### 3.2 Estimating transition probabilities between cell types

**Equation (5)** has given the conditional probability (*p*_*β* →*α*_) that the random walker moves from cell *β* to cell *α*, whereas **Eq. (7)** has given the stationary visit probability of cell *α*, i.e., *p*_*α*_. On the basis of these, *Topographer* estimates the transition probability at which a random walker visits the *j*th cell type from the *i*th cell type (denoted by *q*_*i*↷*j*_), according to

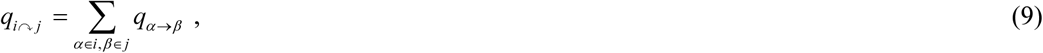

and the transition probability at which the random walker exits the *i*th cell type (denoted by *q*_*i*↷_), according to

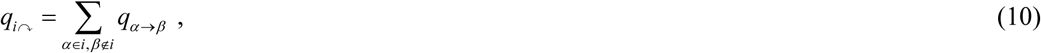

where the unrecorded visit rate on a link, *q*_*β* →*α*_ is given by

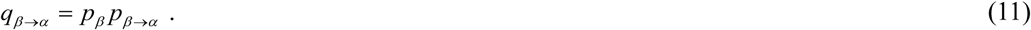

### 3.3 Estimating fate probabilities of cell types

The fate probability for cell type *i*, denoted by *fate*_*i*_, is defined as

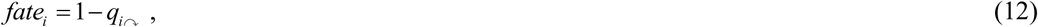

which implies that a larger transition probability at which the random walker exits cell type *i* corresponds to a smaller fate probability for this cell type. This definition is in accordance with our intuition, so it is reasonable. Again, we emphasize that the above formulae for transition probability (*q*_*i*↷*j*_) and fate probability (*fate*_*i*_) have all made use of the pseudotime information.

### 4. The network module infers marker gene networks and their pseudo-temporal changes

In a complex mixture of cells, correlations of gene expression patterns would arise from differences between different cell lineages. To explore the correlation between the patterns of gene expression across development, *Topographer* constructs a series of gene regulatory networks (GRNs) along the pseudotime, which are directed networks for gene-gene interactions. Unsupervised GRNs are then created by GENIE3^21^ that takes advantage of the random forest machine learning algorithm.

Based on the constructed GRNs, *Topographer* further explores the covariation partners of some particular gene (or genes) using a topological network analysis scheme^20^. The method is to identify the set of those genes that are most closely correlated with a given gene (or genes) of interest and that most closely correlate to each other, at a given pseudotime point (in practical calculations, we use data at a pseudotime window to improve accuracy) (in **Fig. 4** of the main text, however, we showed how neighborhood networks of a marker gene change at several representative pseudotime points). **Supplementary Information** provides more details of the method.

### 5. The burst module infers pseudo-temporal characteristics of transcriptional bursting kinetics

Transcriptional bursting kinetics can be characterized by burst size and burst frequency. As is well known, Gamma distributions can well capture this bursty expression in some cases. *Topograph*er uses a Gamma distribution to infer dynamic characteristics of transcriptional bursting kinetics along the cell-state transition trajectories identified from single-cell RNA-seq data. Assume that this distribution takes the form^24^

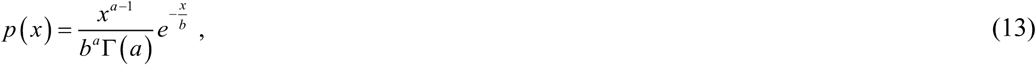

where *x* represents the number of transcripts, *a* represents the mean burst frequency (i.e., the mean number of mRNA production bursts per cell cycle) whereas *b* does the mean burst size (i.e., the average size of the mRNA bursts), and Γ(·) is the common Gamma function.

Thus, in order to infer pseudo-temporal characteristics of transcriptional bursting dynamics, the key is to estimate two parameters *a* and *b* from the dataset at every pseudotime point. For this, *Topographer* makes use of the maximum likelihood method^26^. Since the number of cells at a single pseudotime point would be very few, *Topographer* uses the cell data in a window of this point to obtain more reliable estimations of *a* and *b*.

### 6. Data availability and software

Single-cell data on development of primary human myoblasts can be downloaded from doi:10.1038/nbt.2859^12^. A MATLAB package for the *Topographer* algorithm is available through github (https://github.com/cellfate/topographer).

