## Supplementary Materials for "Topographer Reveals Dynamic Mechanisms of Cell Fate Decisions from Single-Cell Transcriptomic Data"

#### Supplementary Information includes

**1. Motivation**

**2. Data pre-processing**

2.1 Cleaning data

2.2 Identifying the genes of highly differentiating expression

2.3 Calculating distance matrix

3. Details of Topographer

3.1 The backbone module: Identifying the backbone of cell trajectories

3.1.1 Calculating local pseudo-potential

3.1.2 Periphery cells (edge points or edge cells)

3.1.3 Finding valley floors

3.1.4 Setting step length

3.1.5 Trajectory refinement

3.1.6 Pseudotime assignment

3.2 The landscape module: Constructing a quantitative Waddington's landscape

3.2.1 Calculation of cell potential

3.2.2 Scatter plot of developmental landscape

3.2.3 Simple discussion on potential landscape

3.3 The dynamics module: Estimating fate and transition probabilities

3.3.1 Determining cell types

3.3.2 Estimating transition probabilities between cell types

3.3.3 Estimating fate probabilities of cell types

3.3.4 Cell population dynamics: Analysis of a simple example

3.4 The network module: Inferring marker gene networks and their pseudotemporal changes

3.5 The burst module: inferring pseudotemporal characteristics of transcriptional bursting kinetics

3.6 GO enrichment analysis

3.7 Figures for demonstrating *Topographer*'s robustness against the noise in a dataset

**4. Analysis of a toy gene model**

**5. Result demonstrations for analysis of real data examples**

5.1 The development of somatic stem cells

5.2 The differentiation of hematopoietic stem cells

### 1. Motivation

Cell differentiation or development is a complex process, involving multiple cell fate decision points. This process can figuratively be represented by a hierarchical tree in which the multipotent stem and progenitor cells are at the root whereas the mature differentiated cell types are at the bottom with various possible precursor cells representing intermediate cell states. Previous works studied, based on omics data available, cell fate decisions separately from the views of cell lineage committing dynamics (macroscopic, referring to **Supplementary Fig. 1A**), dynamics of gene regulatory networks (mesoscopic, referring to **Supplementary Fig. 1B**), and transcriptional bursting kinetics of genes (microscopic, referring to **Supplementary Fig. 1C**). Significant progresses have been made[1-5], but because these three different scales are coupled and interplayed, the independent studies cannot well elucidate developmental pathways nor can correctly reveal the essential mechanisms of cell fate decisions. This is mainly due to the limitations of omics data themselves.

Recent single-cell measurement technologies such as single-cell RNA-seq, PCR, and mass cytometry, which are enabling generation of data with high resolution, offer a great opportunity to elucidate both developmental processes and cell fate decisions, but require computational algorithms capable of exploiting this resolution. Although previously developed algorithms[1,2,6-16] can successively position single cells in some (not all) single-cell data, this pseudo-temporal positioning is only the first step towards understanding developmental processes and dissecting cell fate decisions. Many important yet fundamental issues, e.g., in single-cell transcriptome data representing a complete development process, e.g., how many cell types there are, how cell types are identified, how one cell type transitions another, and how fate transitions are related to gene networks and transcriptional bursting, remain unsolved. Using high-dimensional single-cell transcriptome data, *Topographer* aims to reveal the integrative mechanisms of cell fate decisions in a coherent way at three scales: cell population dynamics, gene networks dynamics, and transcriptional bursting kinetics, referring to **Supplementary Fig. 1**. In brevity, *Topographer* aims to solve two classes of issues related to cell fate decisions based on single-cell data: 'how' and 'why'.

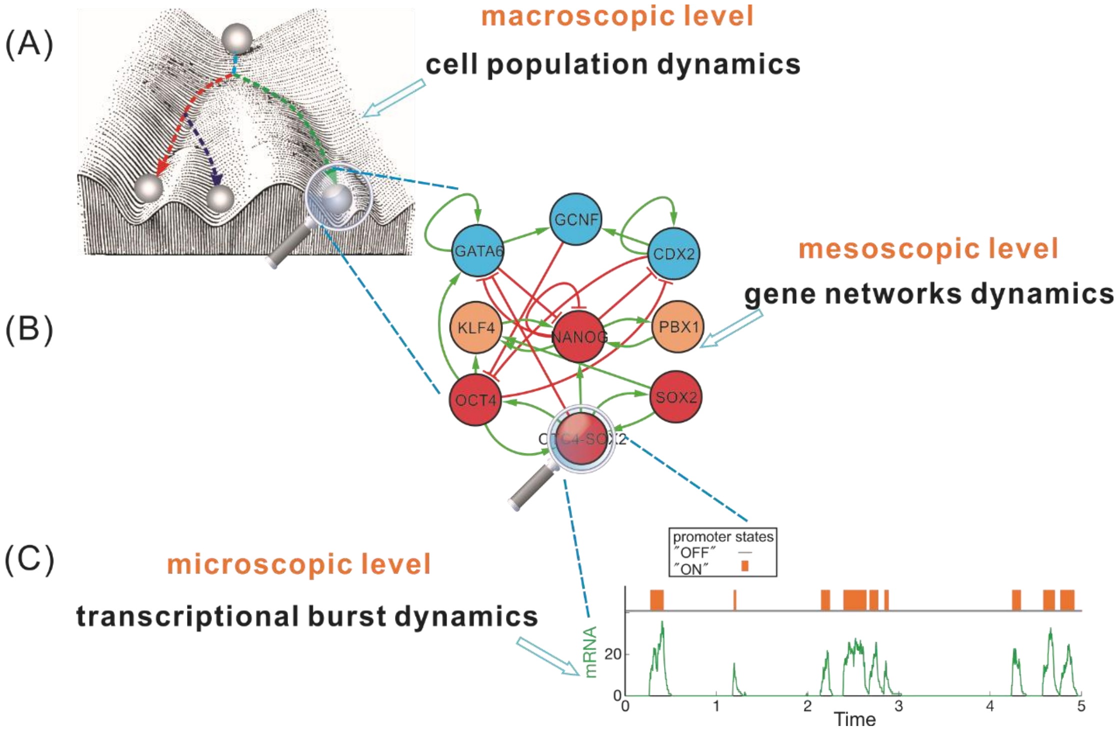


**Supplementary Figure 1** A schematic diagram for study of cell fate decisions involving three different levels. Traditional works separately studied cell population dynamics (A), gene networks dynamics (B), and transcriptional bursting kinetics (C). We aim to reveal the integrative mechanism of cell fate decisions in a coherent way at these three scales, based on singe-cell RNA-seq data. For this, we develop a bioinformatics pipeline, termed *Topographer*, which not only can construct a quantitative (i.e., every cell is loaded with spatiotemporal information) developmental landscape and reveal stochastic dynamics of cell types, but also can infer both marker gene networks and their pseudo-temporal changes as well as dynamic characteristics of transcriptional bursting kinetics across development.

*Topographer* is a bioinformatic pipeline, comprising five functional modules (referring to **Supplementary Fig. 2**): (1) identifying the backbone of cell trajectories from single-cell transcriptomic data; (2) constructing a quantitative developmental landscape of singe-cell data; (3) estimating stochastic cell-type dynamics from single-cell data; (4) inferring marker gene networks and their dynamic changes along the cell-state transition trajectories; and (5) inferring dynamic characteristics of transcriptional bursting kinetics along developmental trajectories. A complete description of *Topographer* is given below.

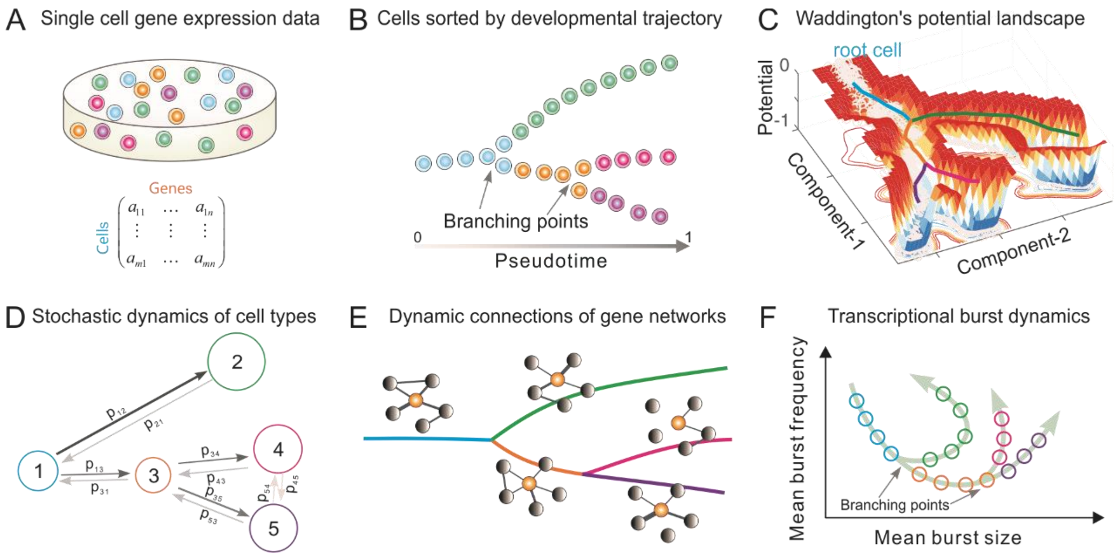


**Supplementary Figure 2** Overview of *Topographer*. The overall *Topographer* comprises five functional modules: (1) the backbone module (**B**); (2) the landscape module (**C**); (3) the dynamics module (**D**); (4) the network module (**E**); and (5) the burst module (**F**). In (A)-(F), the same color represents the same type cell, and the only (C) subfigure is not schematic but is plotted using a set of data generated by a toy model.

(A) Single-cell data are represented by a matrix, where different colors represent different types of cells.

(B) The backbone module identifies the backbone of cell trajectories from the data, where the full pseudotime is set as the interval between 0 and 1.

(C) The landscape module constructs an intuitive Waddington's potential landscape where every cell is equipped with both potential and pseudotime.

(D) The dynamics module reveals stochastic dynamics of cell types by estimating the transition probabilities (indicated by $p_{ij}$ ) among cell types and their fate probabilities, where numbers 1~5 represent cell types, the size of circle represents that of fate probability, and the thickness of arrow represents the size of transition probability.

(E) The network module infers marker gene networks and their dynamic changes along the identified cell trajectories, where the orange ball represents the marker gene, and the thickness of connection line represents the strength of correlation.

(F) The burst module infers dynamic characteristics of transcriptional bursting kinetics (characterized by both burst size and burst frequency) along the pseudotime, where arrows represent the pseudotime direction.

#### 2. Data pre-processing

*2.1 Cleaning data*

Assume that there are $m$ cells and $n$ genes in single-cell RNA-seq data of interest, which are represented as $m$ points in the $n$ -dimensional space ( $X \subseteq \mathbb{R}^n$ ) of gene expression (called the cell state space for convenience). The data for transcript counts for all the genes in all the cells in the dataset are stored in a matrix, $G = \left(g_{ij}\right)_{m\times n}$, where every row is a cell state vector in the cell state space. *Topographer* will be established based on this input matrix.

The first step of data pre-processing is to remove the cells with all low transcript counts. It is needed that the dataset contains at least 1,000 transcripts per cell. The second step is to filter out lowly expressed genes: the genes expressed with less than three transcripts in at least a single cell are discarded.

*2.2 Identifying genes of highly differentiating expression*

First, a function is used to calculate the average and dispersion of every gene in the dataset. Second, the genes in the dataset are first divided into a number of bins based on their average expressions, and z-scores for dispersion within every bin are then calculated. The purpose of this treatment is to identify the genes of highly differentiating expression while controlling the strong relationship between variability and average expression. **Supplementary Fig. 3** demonstrates the result obtained by analyzing a data example.

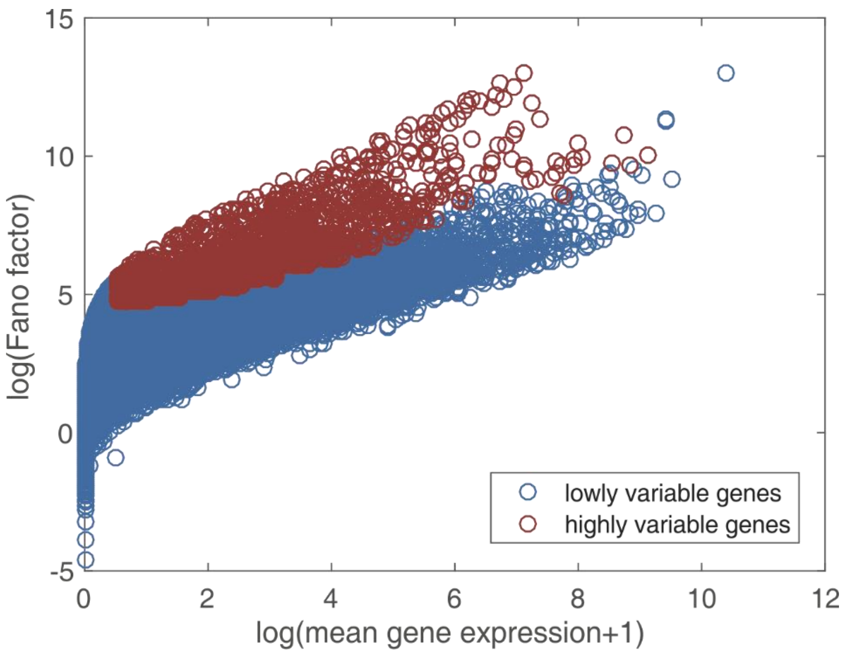


**Supplementary Figure 3** Shown are the differentiating expressions of genes in a dataset in a logarithmic plane of Fano factor and mean expression level adding 1, where Fano factor is defined as the ratio of variance over mean.

*2.3 Calculating the distance matrix*

Let $\boldsymbol{x}$ and $\boldsymbol{y}$ represent the states of the $i$th and $j$th cells in the cell state space (i.e., represent the $i$th and $j$th rows of matrix $G$), respectively. *Topographer* adopts the distance defined by

$$d(\boldsymbol{x},\boldsymbol{y}) = \sqrt{\sum_{k=1}^{n}\left(g_{xk} - g_{yk}\right)^2}, \quad (1)$$

which represents the Euclidean distance between two cell states. *Topographer* may also adopt other kinds of distances such as the Pearson correlation distance.

#### 3. Details of *Topographer*

The overall *Topographer* comprises five functional modules: (1) the backbone module that identifies the backbone of cell trajectories; (2) the landscape module that constructs a quantitative developmental landscape; (3) the dynamics module that estimates stochastic cell-type dynamics characterized by fate and transition probabilities; (4) the network module that infers marker gene networks and their dynamic connections along cell trajectories ; and (5) the burst module that infers dynamic characteristics of transcriptional bursting kinetics along developmental trajectories. Below, we describe each functional module separately.

#### 3.1 The backbone module: identifying the backbone of cell trajectories

##### *3.1.1 Calculating local pseudo-potential*

First, a local density for every cell in the dataset is calculated by applying a Gaussian filter on cell states scattered in the cell state space. That is, for cell state $\boldsymbol{x}$ in $X \subseteq \mathbb{R}^n$, its local density is defined as

$$\rho(\boldsymbol{x}) = \sum_{\boldsymbol{y}\in X} \exp\left(-\frac{d(\boldsymbol{x},\boldsymbol{y})^2}{2\sigma^2}\right), \tag{2}$$

where $d$ is the Euclidean distance between two cell states, $\boldsymbol{x}$ and $\boldsymbol{y}$, seeing to **Eq. (1)**. Thus, a local cell density mostly accounts for the number of cells in the neighborhood defined by $\sigma$. In this implementation, *Topographer* allows users to specify $\sigma$ by setting a parameter $r_{quantile}$ (characterizing the sparseness of the dataset), and then sets $\sigma$ to be the corresponding quantile for all pair-wise distances of cell states in $X$.

Then, based on local density function $\rho(\boldsymbol{x})$ for cell state $\boldsymbol{x}$, the pseudo-potential of this cell is defined as

$$\tilde{E}(\boldsymbol{x}) = -\log\rho(\boldsymbol{x}) \ . \tag{3}$$

In the following, local density and local pseudo-potential will be alternatively used due to **Eq. (3)** unless confusion arises.

##### *3.1.2 Periphery cells (edge points or edge cells)*

Before finding a valley-floor trajectory in state space $\mathbb{R}^n$, it is needed to mark out those cells that fall around the peripheral region of the density distribution in the cell state space. Such periphery cells are then excluded from the candidate of pseudo-potential valley in order to improve accuracy. A cell, $\boldsymbol{x}$, is marked as periphery if its local density is relatively lower in its neighborhood defined as $U(\boldsymbol{x}) = \{\boldsymbol{y} | d(\boldsymbol{x},\boldsymbol{y}) \le \sigma\}$, that is, if

$$\rho(\boldsymbol{x}) = Q_r\left(\{\rho(\boldsymbol{y}) | \boldsymbol{y} \in U(\boldsymbol{x})\}\right) \ , \tag{4}$$

where $Q_r(u)$ is a quantile of the discrete set of values $u$, and $r = 0.2$ may be set.

##### *3.1.3 Finding valley floors*

Possible branching trajectories representing valley floor lines of a landscape in the dataset can be constructed through a recursive process. In this process, a method that has analogy with the density-based clustering method developed originally by Rodriguez et al.[17] is adopted to find valleys with local minimal pseudo-potentials on the perimeter region centered at some cell state.

In order to search for a pseudo-potential valley trajectory in the dataset $X$, *Topographer* finds local minimal pseudo-potential valley floors around the starting point selected according to the prior knowledge. By connecting the corresponding cell states to the starting point, the resulting segments are considered as one part of a pseudo-potential valley floor. For the rest part, the same procedure is repeated recursively with the newly found valley points as orbital centers, thus finding all pseudo-potential valley floors on the corresponding orbits (referring to **Supplementary Fig. 8-10**).

It should be pointed out that for an orbital center other than the starting point, the found valley floors would include a point on the "reversed direction", which is not expected in our algorithm. To handle such an exception, *Topographer* excludes those valleys that are too close to a previous cell state in the valley floor, that is, *Topographer* excludes those points in distances smaller than the step lengths. Also, any two newly found valley floors in the distance smaller than the step length would be merged by abandoning valleys with larger pseudo-potentials (or lower local densities) (referring to **Supplementary Fig. 8-10**). Such a treatment may greatly improve the algorithm's robustness against the noise in the dataset.

Now, let us state details of finding valley floors. First, the search method, i.e., a method of finding pseudo-potential wells on circular tubes (every such a tube is below called a super ring for convenience) in the cell state space, which is analogous to a density-based approach developed originally by Rodriguez and colleagues[17], is based on the idea that cluster centers are characterized by a lower pseudo-potential than their neighbors and by a relatively larger distance from any points with lower pseudo-potentials. This idea forms the basis of a procedure to find pseudo-potential wells on a super ring. In this procedure, the number of pseudo-potential wells arises intuitively, outliers are automatically spotted and excluded from extra analysis, and pseudo-potential wells are recognized regardless of their shape and of the dimensionality of the space in which they are embedded. Then, if we define

$$\delta(\boldsymbol{x}) = \begin{cases} \max_{\boldsymbol{y}\neq\boldsymbol{x}} d(\boldsymbol{x},\boldsymbol{y}) & \text{if } \tilde{E}(\boldsymbol{x}) = \min_{\boldsymbol{y}\in X} \tilde{E}(\boldsymbol{y}) \\ \min_{\boldsymbol{y}:\tilde{E}(\boldsymbol{y})<\tilde{E}(\boldsymbol{x})} d(\boldsymbol{x},\boldsymbol{y}) & \text{otherwise ,} \end{cases} \quad (5)$$

then finding pseudo-potential wells on the super ring is based on a combination of relatively smaller $\tilde{E}$ and relatively larger $\delta$. For convenience, this approach is called as the pseudo-potential well search algorithm. If the given single-cell data are sufficiently smooth, then the segments linking the center and the found potential wells on the super ring can be taken as approximate parts of the entire developmental trajectory.

To find out local pseudo-potential valleys in the dataset, *Topographer* uses a combination of relatively larger $\rho$ (or relatively smaller $\tilde{E}$) and relatively larger $\delta$. That is, after setting a parameter $k$, the data point $\boldsymbol{x}_k$ with the $k$-th largest $\gamma$ (which is equal to the product of $\rho$ and $\delta$, i.e., $\gamma = \rho\delta$) is firstly chosen. Any data point with larger $\rho$ (or smaller $\tilde{E}$) and larger $\delta$, that is, any $\boldsymbol{x}$ satisfying $\rho(\boldsymbol{x}) \geq \rho(\boldsymbol{x}_k)$ (or $\tilde{E}(\boldsymbol{x}) \leq \tilde{E}(\boldsymbol{x}_k)$) and $\delta(\boldsymbol{x}) \geq \delta(\boldsymbol{x}_k)$, is used to define a local pseudo-potential valley.

A complete valley floor is constructed by terminating the recursive process at orbital centers where there are no local pseudo-potential valleys around them. Also, since no loops are assumed to exist in the developmental trajectory, any local pseudo-potential valleys that are too close to the found trajectory segments should be abandoned. This may also evoke termination of the recursive search by the rule described above.

##### *3.1.4 Setting step length*

Since every segment is constructed by connecting an orbital center with a valley floor found on the orbital region, the orbit radius determines a step length in searching for pseudo-potential valley floors. There are two approaches to set the step length: One is to calculate a fixed radius for all cell states in the dataset, and the other is to calculate a specific step length for every cell state found on the pseudo-potential valley.

*Topographer* uses a cell state-dependent step length in calculation. The merit is that it can make the algorithm be optimally robust in analysis of dataset with highly heterogeneous, local pseudo-potential valleys. For example, in a region in the state space where the local pseudo-potential of cell state is low in the dataset, the neighbors of any local pseudo-potential valleys would be so sparse that a long step is desired to compensate for the error induced by noise in the dataset. On contrary, shorter steps would be more advantageous in regions with lower pseudo-potentials, providing a better resolution and therefore increasing the accuracy in finding a valley floor. *Topographer* defines a cell state-dependent step length to achieve these features, that is

$$s(\boldsymbol{x}) = \alpha \min\left\{ d(\boldsymbol{x}, \boldsymbol{y}) \middle| \rho(\boldsymbol{y}) \le \frac{1}{\sqrt{e}} \rho(\boldsymbol{x}), \boldsymbol{y} \in X \right\} , \tag{5}$$

where $\boldsymbol{x}$ is a cell state at the pseudo-potential valley and $\alpha$ is a factor. Such a step length resembles an estimation of the standard deviation of a normally distributed set of data values, since the probability density function $f(\boldsymbol{x}) = \frac{1}{\sqrt{2\pi}\sigma} e^{-d(\boldsymbol{x},\boldsymbol{0})/(2\sigma^2)}$ decreases to $\frac{1}{\sqrt{e}}$ times of its peak value at $|\boldsymbol{x}| = \sigma$. Thus, multiplying it with a factor $\alpha$ (which is set as 1.5 in our cases) may provide a good estimation of the "width" of the trajectory in the corresponding region.

##### *3.1.5 Trajectory refinement*

After the above search procedure is finished, the found pseudo-potential valleys are stored in an undirected acyclic graph (a tree). To achieve better accuracy and coverage, the pseudo-potential valley tree is refined by searching for an intermediate cell state with low local potential for each floor, and extending each ending segments to the edge cells. Then, the user is required to select a cell as the root cell state of a developmental trajectory. With this cell state taken as a root, the developmental tree is thus constructed, which is actually a directed acyclic graph. Additionally, all the branches can be calculated by finding the paths from the root node to each leaf node in the developmental tree. Each path, represented by a sequence of cell states, accounts for a single branch in the data, while any two of such paths may share a common sequence of cell states in the early parts.

In particular, when the selected root cell state is not a leaf node of the tree, it is needed to extend the segments connected to this state in all the ‘reversed’ directions at the first step (i.e., the step of finding pseudo-potential wells on a super ring). To be exact, *Topographer* finds the inter-set of the extending region of those segments and picks up a cell state with the locally lowest potential, which is connected to the root cell state. After this, the same procedure of trajectory refining and root selecting is again carried out to utilize the extended version of the pseudo-potential valley tree.

##### *3.1.6 Pseudotime assignment*

To completely order the captured cell states on a developmental trajectory, it is needed to assign a pseudotime for each cell in the cell state space. Before that, a root node in the tree is first selected based on the rule that the sum of the distances to the initial cells that the user provides or sets is smallest. Thus, this root node corresponds to the shortest distance to the initial cells, and is denoted as $x_{progen}$. Then, every cell state is assigned to a tree node with the shortest Euclidean distance. At this procedure, each cell state is also assigned to a unique branch in the developmental trajectory.

Specifically, an integer pseudotime is initially assigned to every cell state in the developmental tree according to the geodesic distance to the root node $x_{progen}$. For the rest of cell states, a pseudotime is assigned to a cell state by projecting on a closest edge (i.e., the edge linking the nearest node to the secondary nearest node) on the tree. More precisely, every cell state is perpendicularly projected onto the closest edge connected to its related node. A pseudotime is then obtained by averaging the pseudotime of the two end nodes weighted by the Euclidean distance.

**Supplementary Fig. 4** shows a flowchart for how the backbone of a developmental trajectory is found in the multi-dimensional cell state space. More details are shown in **Supplementary Fig. 8-10**.

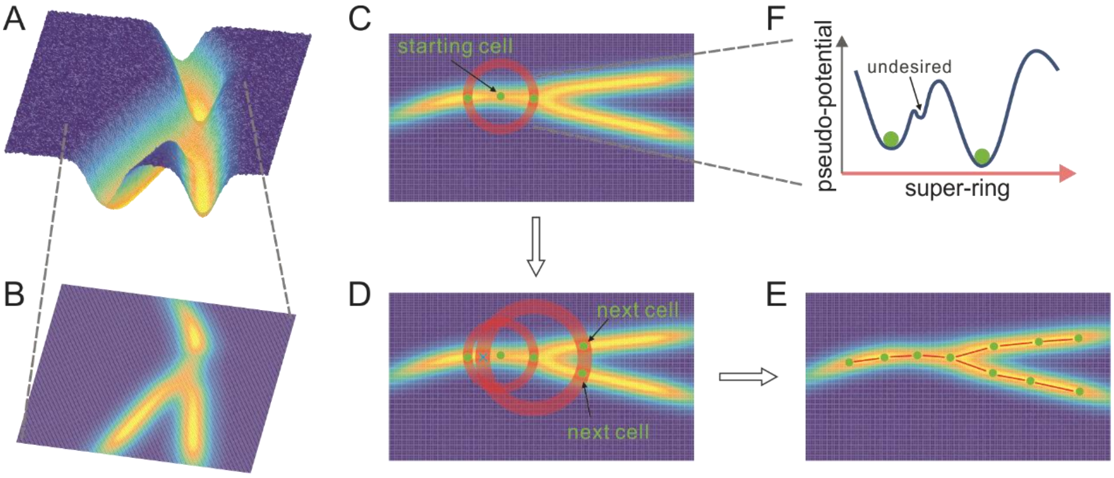


**Supplementary Figure 4** A Schematic diagram for an algorithm to find the backbone of cell trajectories in a multi-dimensional cell state space, where (A) shows a branched trajectory in a high-dimensional space whereas (B) shows a projection of this trajectory, **(C)-(E)** show a flowchart (indicated by arrows): First, *Topographer* selects an initial cell **(C)** as the center of a super-ring in the cell state space and searches for pseudo-potential wells on this ring. Then, *Topographer* repeats recursively on every newly found pseudo-potential well **(D)** until no pseudo-potential wells are found, thus obtaining a tree-like backbone of cell trajectories **(E)**. Finally, *Topographer* projects every cell onto the backbone, thus ordering all the cells in the dataset. **(F)** shows a super-ring example, where two undesired pseudo-potential wells and an undesired pseudo-potential well are indicated.

#### 3.2 The landscape module: constructing a quantitative Waddington's landscape

##### *3.2.1 Calculation of cell potential*

After the backbone of a developmental trajectory has been identified and every cell has been equipped with a pseudotime, the landscape module is to construct a developmental landscape. For this, *Topographer* calculates a potential of every cell in the dataset (see the definition below). It is expected that the potential to be introduced can be avoid the shortcomings of the pseudo-potential, e.g., it cannot correctly reflect transition probabilities among different cell types since there would be probability fluxes between cells due to cell division, cell death, and so on. In this calculation, *Topographer* analogizes transitions between cells at distinct stages of the developmental process to a random walker who moves randomly between the data points scattered in the cell state space[18].

In order to construct a weighted directed graph $\mathbf{W}$, the key question to be solved is how *Topographer* uses the pseudotime information. Assume that the cells for one branching trajectory progress along the pseudotime in a linear manner. If the weight of the directed edge from cell $\alpha$ to cell $\beta$ is denoted by $W_{\alpha\to\beta}(\tau)$, where $\tau$ represents the pseudotime, then the assumption implies the linear ordinary differential equation

$$\frac{d}{d\tau}W_{\alpha\to\beta}(\tau) = -\chi W_{\alpha\to\beta}(\tau) \ , \tag{6}$$

where $\chi$ is a positive constant, representing the rate that cell $\alpha$ transitions to cell $\beta$. If the initial condition is denoted by $W_0$, then the solution to **Eq. (6)** can be expressed as

$$W_{\alpha\to\beta} = W_0 e^{-\chi(\tau_\alpha - \tau_\beta)} \ , \tag{7}$$

where $\tau_\alpha$ and $\tau_\beta$ represents the pseudotime points corresponding respectively to cells $\alpha$ and $\beta$. Thus, $\chi$ can be viewed as a scaling factor to adjust the weight difference between forward and backward directions. The setting of $\chi$ in general depends on the dataset of interest but the value of $\chi$ is set as 30 in our situations. Apparently, the larger the value of $\chi$ is set, the larger is the weight along the positive direction of the pseudotime. On contrary, the weight is smaller. **Eq. (7)** indicates that the weight of the directed edge from a cell to another changes with the pseudotime in an exponential manner. We stress that the weight given by **Eq. (7)** has used the information on the identified pseudo-temporal cell trajectories.

Then, in order to determine cell visit probability on a random walk, *Topographer* defines a conditional probability that the random walker moves from cell $\beta$ to cell $\alpha$ as the relative link weight, given by

$$p_{\beta\to\alpha} = \frac{W_{\beta\to\alpha}}{\sum_{\beta} W_{\beta\to\alpha}} \quad , \tag{8}$$

It is easily seen that $p_{\beta\to\alpha}$ is independent of $W_0$. Denote by $p_\alpha$ the stationary visit probability of cell $\alpha$. Then, this probability can in principle be derived from a recursive system of the form

$$p_\alpha = \sum_{\beta} p_\beta p_{\beta\to\alpha} \tag{9}$$

Therefore, $p_\alpha$ represents the probability that the random walker visits the $\alpha$ cell from all the other cells. Note that **Eq. (9)** is actually a master equation[19] and can efficiently be solved with the power-iteration method[20]. However, to ensure that a unique solution of this equation is independent of the starting node in the directed network, the random walker instead teleports to a random node at a small rate $\tau$. Furthermore, to obtain more robust results depending less on the teleportation parameter $\tau$, it is most often to use teleportation to a node proportional to the total weight of the links to the node[18]. With these considerations, the stationary cell visit probability is then modified as

$$p_\alpha = (1-\tau)\sum_{\beta} p_\beta p_{\beta\to\alpha} + \tau \frac{\sum_{\beta} W_{\alpha\to\beta}}{\sum_{\alpha,\beta} W_{\beta\to\alpha}} \quad , \tag{10}$$

Finally, *Topographer* estimates the potential of every cell in the dataset according to

$$E_\alpha = -\log p_\alpha \quad , \tag{11}$$

where $\alpha$, an index, represents a cell in the dataset. Apparently, the potential defined in such a manner has used the information on the identified cell trajectories due to **Eq. (7)**, but pseudo-potential does not consider pseudotime (in fact, it unnecessarily considers pseudotime since pseudo-potential is defined using cell density), referring to the difference between pseudo-potential (**Fig. 5A** and **5B**) and potential (**Supplementary Fig. 5C** and **5D**). After every cell is equipped with a potential, all these potentials are then used to construct a potential landscape.

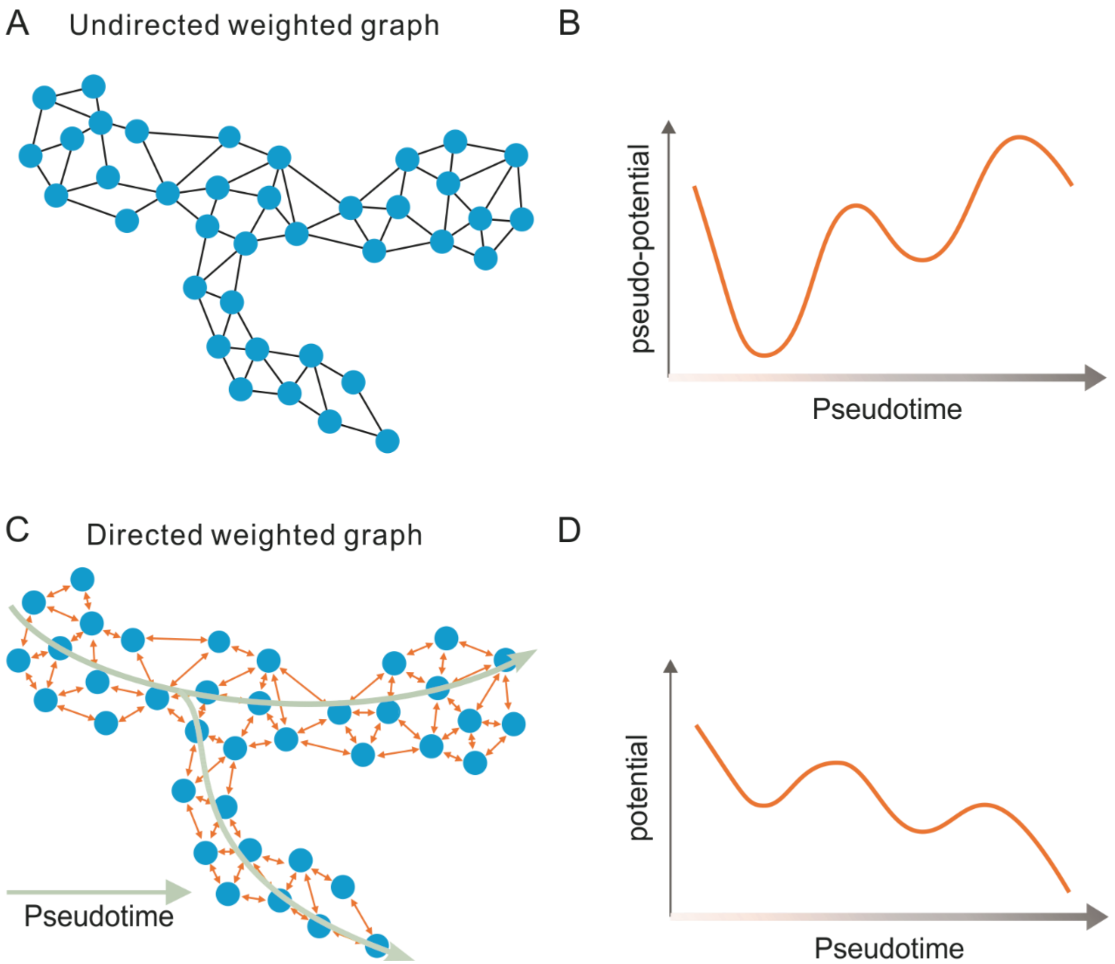


**Supplementary Figure 5** Schematic diagram for the difference between potential and pseudo-potential: (A, B) the undirected weighted graph and the corresponding pseudo-potential along the pseudotime; (C, D) the directed weighted graph and the corresponding potential also along the pseudotime. We point out that: (1) the connection between cells is endowed with the pseudotime in (C) but not in (A); (2) in a Waddington's potential landscape, a 'ball' moves from a higher potential to a lower potential. Thus, the potential rather than the pseudo-potential is in accord with our intuition.

##### *3.2.2 Scatter plot of developmental landscape*

After all the cells are equipped with potentials, all these potentials are then used to draw a potential landscape. Here we introduce two methods for such a drawing.

The first method is stated as follows. First, the tSNE method[21] or the PCA method[22] is used to reduce dimensions for visualization. In general, dimension reduction cannot well reflect the information on coordinates in the visualized landscape, e.g., PCA1 and PCA2 in **Fig. 3C** in the main text and **Supplementary Fig. 10A** below do not represent concrete components in the dataset.

Second, *Topographer* uses the nearest neighbor interpolation method to perform interpolation on a 2- or 3- dimensional scattered data set. Specifically, *Topographer* uses ScatteredInterpolant (a function of the MATLAB software) to establish corresponding relationships between a set of points, $(x, y)$, and a set of the modified cell potentials given by **Eq. (10)**, $E$. These corresponding relationships, denoted by $E = F(x, y)$, in principle define a curved surface in the 3-dimensional space of potential landscape, which in return passes through all the sampling points in the underlying space. After that, *Topographer* uses the nearest neighbor interpolation to evaluate this surface at any query point, $(x_q, y_q)$, thus obtaining an interpolating value of every known potential given by **Eq. (11)**, i.e., $E_q = F(x_q, y_q)$. Third, a Gaussian kernel is used to smooth interpolation, and the identified developmental trajectory is drawn on the landscape (referring to the thick green lines in **Supplementary Fig. 6A** and **6C**). To that end, the drawing of a 'stereometric' developmental landscape is finished (referring to **Supplementary Fig. 6** and **13A** later).

The second method is stated below. Similar to the first method, the tSNE method[21] or the PCA method[22] is used to reduce dimensions for visualization, but the scatter plot of a developmental landscape needs to fit a curved surface in a dimension-reduced space. The method is as follows: for every pseudotime point $\tau$, define

$$U_\tau(x) = -\ln p_\tau(x) \tag{12}$$

where

$$p_\tau(x) = \sum_{k=1}^{K} \frac{a_k}{\sqrt{2\pi}\sigma_k} e^{-(x-\mu_k)^2/(2\sigma_k^2)} \tag{12a}$$

is a hybrid Gaussian distribution with nonnegative parameters, $a_k$ satisfies the constrained condition $\sum_{k=1}^{K} a_k = 1$. In **Eq. (12a)**, $K$ represents the number of branches, e.g., a single branching trajectory corresponds to $K = 1$ whereas a bi-branching trajectory to $K = 2$. Parameters $a_k$, $\sigma_k$, and $\mu_k$ can be obtained by fitting the data (i.e., by fitting the known potentials for the cells at pseudotime point $\tau$). Then, the obtained $U_\tau(x)$ gives a curve in the dimension-reduced space. Furthermore, the split joint of all these curves along the pseudotime constitutes a 'stereometric' developmental landscape. **Supplementary Fig. 6A** and **6C** show two Waddington's potential landscapes constructed using an artificial set of data and a set of single-cell data on the differentiation of primary human myoblasts, respecitvely.

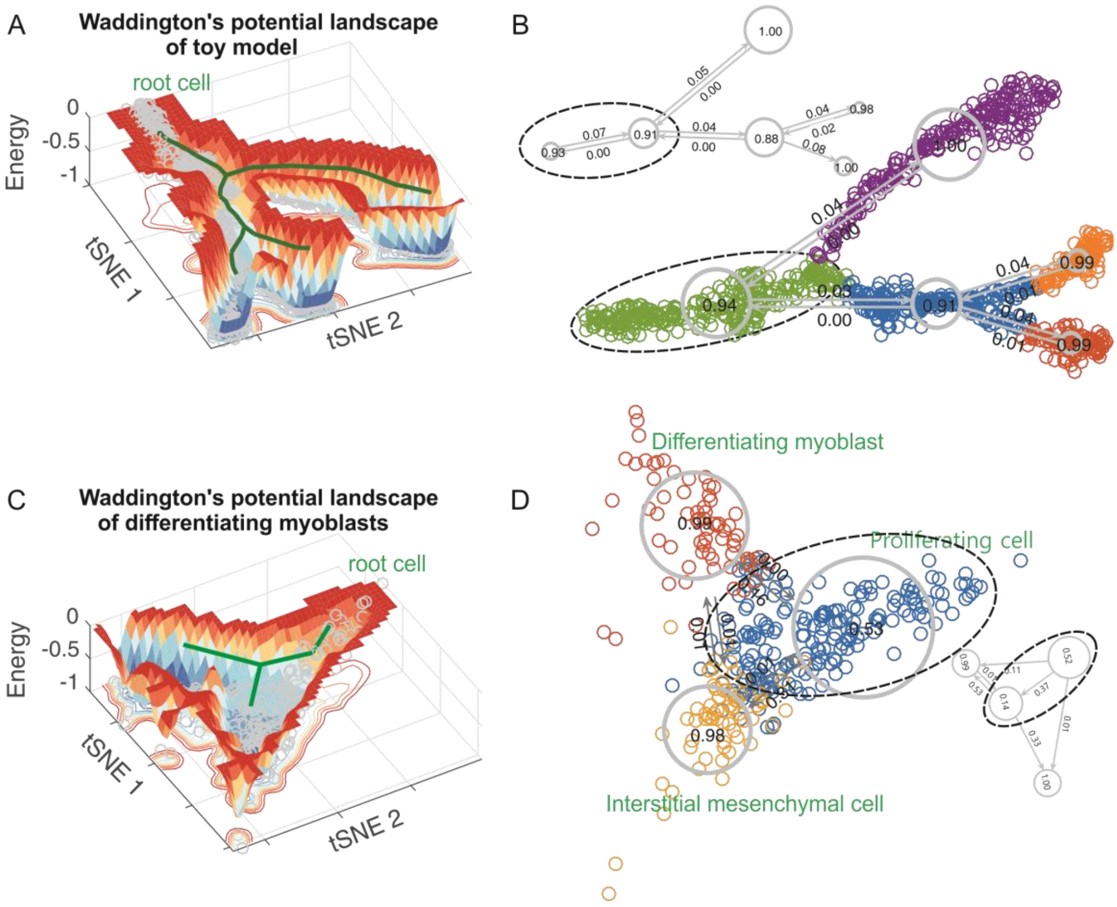


**Supplementary Figure 6.** *Topographer* constructs Waddington's potential landscapes (the first column) and reveals stochastic dynamics of cell types (the second column).

**(A)** A Waddington's potential landscape constructed from an artificial set of data, where a thick green line with branches represents the backbone of cell trajectories, and each small, grey circle represents one cell. tSNE-1 and tSNE-2 are two dummy variables. The normalized potential is shown with depth of color representing the size of potential.

**(B)** Stochastic dynamics of cell types along branching trajectories, revealed from the same set of data as in (**A**), where both the fate probabilities of cell types (distinguished by colors) and transition probabilities among them are indicated. A cell type (below panel) is further divided into two subtypes (top panel), and transition rates are also indicated.

**(C)** The interpretation is similar to **(A)** but the result is obtained by analyzing a set of single-cell data on the differentiation of primary human myoblasts.

**(D)** The interpretation is similar to **(B)** but the result is obtained by analyzing the same set of data as in (**C**). Three known cell types: proliferating cells, differentiating myoblast and interstitial mesenchymal cells, are indicated by dashed ellipse and circles. The dashed ellipse shows that the proliferating cell type (top panel) can further be classified into two subtypes (below panel). The fate probability of every cell subtype and transition probabilities between these subtypes are also indicated.

##### *3.2.3 Simple discussion on potential landscape*

Consider a system of genes and cells across development. Denote by $f(x;\tau)$ the probability that the fraction of the total cell number is $x$, by $g(\boldsymbol{y}|x;\tau)$ the probability that the gene expression levels are $\boldsymbol{y}$ but conditional on $x$, and by $F(x,\boldsymbol{y};\tau)$ the joint probability distribution of $x$ and $\boldsymbol{y}$, at time $\tau$. Then, we can have

$$F(x,\boldsymbol{y};\tau)=f(x;\tau)g(\boldsymbol{y}|x;\tau) \tag{13}$$

Define

$$U_1(x;\tau)=-\ln f(x;\tau),\ \ U_2(\boldsymbol{y}|x;\tau)=-\ln g(\boldsymbol{y}|x;\tau) \tag{14}$$

which represent the potentials of cells and genes at time $\tau$, respectively. Then,

$$U(x,\boldsymbol{y};\tau)=U_1(x;\tau)+U_2(\boldsymbol{y}|x;\tau) \tag{15}$$

represents the potential of the entire system (called a two-level potential) at time $\tau$. Thus, if $U_1(x;\tau)$ or $U_2(\boldsymbol{y}|x;\tau)$ is bimodal, then $U(x,\boldsymbol{y};\tau)$ is not necessarily bimodal, and if the only one of $U_1(x;\tau)$ and $U_2(\boldsymbol{y}|x;\tau)$ is unimodal, then $U(x,\boldsymbol{y};\tau)$ may be bimodal. Traditional studies[9] separately considered $U_1(x;\tau)$ and $U_2(\boldsymbol{y}|x;\tau)$. Apparently, these separate studies cannot correctly describe the characteristics of the potential of the full system. However, the developmental landscapes constructed by *Topographer* (referring to **Supplementary Fig. 6** above and **Supplementary Fig. 13** later) implicitly considered such two-level potentials, and are therefore reasonable. Theoretically describing dynamic characteristics of a developmental landscape needs to establish the so-called balance equation for the gene-cell system, which however is a challenging task.

#### 3.3 The dynamics module: Estimating fate and transition probabilities

##### 3.3.1 *Determining cell types*

In order to quantify cell type dynamics, it is first needed to determine cell types. For this, *Topographer* adopts the following rule: Every branch in the identified developmental trajectory is defined as one kind of cell type, and a different branch is defined as one different kind of cell type. Furthermore, every potential well on every branch is defined as one kind of cell subtype, and a different potential well is defined as one different kind of cell subtype. It should be pointed out that the cell types determined in such a manner are not unique but depend on the choice of $\tilde{E}$ and $\delta$ (see their respective definitions above).

##### 3.3.2 *Estimating transition probabilities between cell types*

**Equation (8)** has given the conditional probability ( $p_{\beta\to\alpha}$ ) that the random walker moves from cell $\beta$ to cell $\alpha$ , whereas **Eq. (10)** has given the stationary visit probability of cell $\alpha$ , i.e., $p_\alpha$ . Then, *Topographer* estimates the transition probability at which a random walker visits the $j$th cell type from the $i$th cell type, according to

$$q_{i\curvearrowright j} = \sum_{\alpha\in i,\beta\in j} q_{\alpha\to\beta} \quad (16)$$

and the transition probability at which the random walker exits the $i$th cell type, according to

$$q_{i\curvearrowright} = \sum_{\alpha\in i,\beta\notin i} q_{\alpha\to\beta} \quad (17)$$

where the unrecorded visit rates on links, $q_{\beta\to\alpha}$ is given by

$$q_{\beta\to\alpha} = p_\beta p_{\beta\to\alpha} \quad (18)$$

##### 3.3.3 *Estimating cell-type fate probabilities*

The fate probability for cell type $i$ , $fate_i$ , is defined as

$$fate_i = 1 - q_{i\curvearrowright} \quad (19)$$

This definition implies that a larger transition rate at which the random walker exits cell type $i$ corresponds to a smaller fate probability for this cell type, which is in accordance with our intuition. Again, we emphasize that the above formulae for transition probability ( $q_{i\curvearrowright j}$ ), and fate probability ( $fate_i$ ) all have used the information on the pseudo-temporal cell trajectories.

**Supplementary Fig. 6B** and **6D** show the fate probabilities of cell types and the transition probabilities between them, obtained according to the above formulae but by analyzing an artificial and a realistic sets of data, respectively.

##### *3.3.4 Cell population dynamics: Analysis of a simple example*

Consider a model of multi-stage cell lineages. This model assumes that *A*-type cells are stochastically divided to *B*- and *C*-type cells with either asymmetric or symmetric cell division, according to the following rule (i.e., a set of chemical reactions)

$$
\begin{aligned}
&\varnothing \xrightarrow{k_0} A \\
&A \xrightarrow{k_1} A + A \\
&A \xrightarrow{k_2 p} B \\
&A \xrightarrow{k_2(1-p)} C \\
&B \xrightarrow{\delta_B} \varnothing \\
&C \xrightarrow{\delta_C} \varnothing
\end{aligned}
\tag{20}
$$

where $k_0$ is the basal production rate for $A$ -type cells, $k_1$ is the proliferation rate whereas $k_2$ is the differentiation rate (we assume $k_2 > k_1$), $\delta_B$ and $\delta_C$ represent the degradation rates at which *B*- and *C*-type cells commit to terminal differentiation and loss, respectively. Parameters $p$ is the probability that *A*-type cells differentiate into *B*-type cells and hence $1-p$ represents the probability that *A*-type cells differentiate into *C*-type cells. Note that parameters $k_1$ and $k_2$ can be estimated from a given set of data using the above method but parameter $\delta_B$ and $\delta_C$ as well as $p$ are in general unknown. Denote by $A(t), B(t), C(t)$ the mean numbers of cells for cell types $A$ , $B$ and $C$ at time $t$ , respectively. Then, the differential equation for the time evolution of the cell populations in every generation reads

$$
\begin{aligned}
\frac{dA(t)}{dt} &= k_0 + (k_1 - k_2) A(t) \\
\frac{dB(t)}{dt} &= k_2 p A(t) - \delta_B B(t) \\
\frac{dC(t)}{dt} &= k_2 (1-p) A(t) - \delta_C C(t)
\end{aligned}
\tag{21}
$$

Note that the time used here is not pseudotime. Solving this linear equation, we get the steady states of the cell numbers for three cell types as follow

$$
\begin{aligned}
A^* &= \frac{k_0}{k_2 - k_1} \\
B^* &= \frac{k_2 p}{\delta_B} \frac{k_0}{k_2 - k_1} \\
C^* &= \frac{k_2 (1-p)}{\delta_C} \frac{k_0}{k_2 - k_1}
\end{aligned}
\tag{22}
$$

From these expressions combined with the definition of pseudo-potential, we can know that if the cell density in $A-$ type cells (similarly in $B-$ and $C-$ type cells) is larger, then the pseudo-potential is smaller. In a Waddington's landscape, however, a 'ball' moves from a higher potential to a lower potential. In addition, pseudo-potential lacks the information on pseudotime (but potential relies on pseudotime). This is a reason why we need to introduce the potential of every cell that is in accord with our intuition when constructing a Waddington's developmental landscape based on single-cell data.

##### 3.4 The network module: inferring mark gene networks and their pseudo-temporal changes

In a complex mixture of cells, the correlations between gene expression patterns could arise from differences between different cell lineages. To explore these regulatory patterns of gene expression across cell development, *Topographer* generates a series of gene regulatory network (GRNs) along the pseudotime, which are the directed networks of gene-gene interactions. Unsupervised GRNs were created using GENIE3[23], an algorithm for the inference of gene regulatory networks from expression data, which decomposes the question of predicting a regulatory network between $m$ genes into $m$ different regression questions.

Based on the constructed GRNs, *Topographer* further explores the covariation partners of some particular gene (or genes) using a topological network analysis scheme. This scheme can identify a set of genes most closely correlated with a given gene (or genes) of interest, and which also most closely correlate with each other[24].

Details of the algorithm are stated as follows

**Inputs**

| | |
|---|---|
| $\mathbf{X} \in \mathbb{R}^{N \times M}$ | Expression matrix with $N$ cells and $M$ genes |
| $g_0$ | "bait" gene index |
| $k$ | Neighborhood size |
| $c$ | Minimal connectivity |

**Algorithm**

1. Construct the distance matrix $\mathbf{D}$ with components $\tilde{d}_{ij} = 1 - \text{Pearson correlation}\left(gene_i, gene_j\right)$, where the correlation is taken over all cells.
2. Construct an unweighted, directed network $\mathbf{G}$ that constitutes the so-called "network

neighborhood of gene $g_0$" using the following method:

(1) Finding the *k*-nearest neighbors of gene $g_0$ and introducing a directed edge from $g_0$ to each of these $k$ genes, with the result stored in set $g_1$.

(2) For every member of set $g_1$, similarly add directed edges to $k$ closest genes, together forming a set $g_2$.

(3) The network is trimmed iteratively by removing any vertex that has fewer than $c$ incoming edges.

**Outputs**

$\mathbf{G} \in \mathbb{R}^{(l+1)\times(l+1)}$, a neighborhood matrix of "bait" gene $g_0$ connected with $l$ genes.

#### 3.5 The burst module: inferring pseudo-temporal characteristics of transcriptional bursting kinetics

Transcription occurs often in a bursty manner and single-cell experimental measurements have also provided evidence for transcriptional bursting both in bacteria and in eukaryotic cells[24]. By analyzing a simplified stochastic model of gene expression, Xie, *et al* showed[26] that the number of mRNAs produced in the bursty fashion followed a Gamma distribution determined by two parameters: one is mean burst frequency (i.e., the mean number of mRNA production bursts per cell cycle), and the other is mean burst size (i.e., the average size of the mRNA bursts).

Transcriptional bursting kinetics can be characterized by burst size and burst frequency. As is well known, Gamma distributions can well capture this bursty expression in some cases. *Topographer* uses a Gamma distribution to infer dynamic characteristics of transcriptional bursting kinetics along the pseudotime from single-cell RNA-seq data. Assume that this distribution takes the form

$$p(x) = \frac{x^{a-1}}{b^a \Gamma(a)} e^{-\frac{x}{b}} \tag{23}$$

where $x$ represents the number of transcripts, $a$ is the mean burst frequency, and $b$ is the mean burst size.

In order to estimate two parameters $a$ and $b$ from the dataset at every pseudo-time point, *Topographer* makes use of the maximum likelihood method[22]. Since the number of cells at a single pseudo-time point would be very few, *Topographer* uses the cell data in a window of this point to obtain more reliable estimations of $a$ and $b$.

##### 3.6 GO enrichment analysis

Before calculation, the genes expressed in levels lower than 5 UMI in more than 95% of the total cell population are excluded. Then, we apply an intuitive method to filter branch-specific genes from the remaining moderately or highly expressed genes. First, a *branch differing score* is assigned to each gene by calculating the Euclidean distance between expression profiles between two branches normalized by the highest expression level. Then, a cutoff value for the branch differing score is set by manual inspection to filter out a set of highly branch-specific genes. The resulting set of genes are classified by the k-medoids algorithm[27] into 2 clusters, with parameter $k = 2$. The results from different $k$ parameter values ranging from 2 to 12 strongly suggested only 2 classes of genes in the branch-differing genes.

A visualized GO enrichment analysis is carried out by ClueGo[28] plug-in with the Cytoscape software to characterize the resulting 2 clusters of branch-specific genes. For each cluster, terms related to any of the clustered genes in the level from 4 to 5 are collected in the GO database with aspect to the underlying biological process and are functionally grouped by ClueGo. Each group is marked by the term with highest significance in the group. Significance (p-value) is calculated with two-sided hypergeometric test regarding the mouse Ontology Reference Set. The results are shown in **Supplementary Fig. 7**.

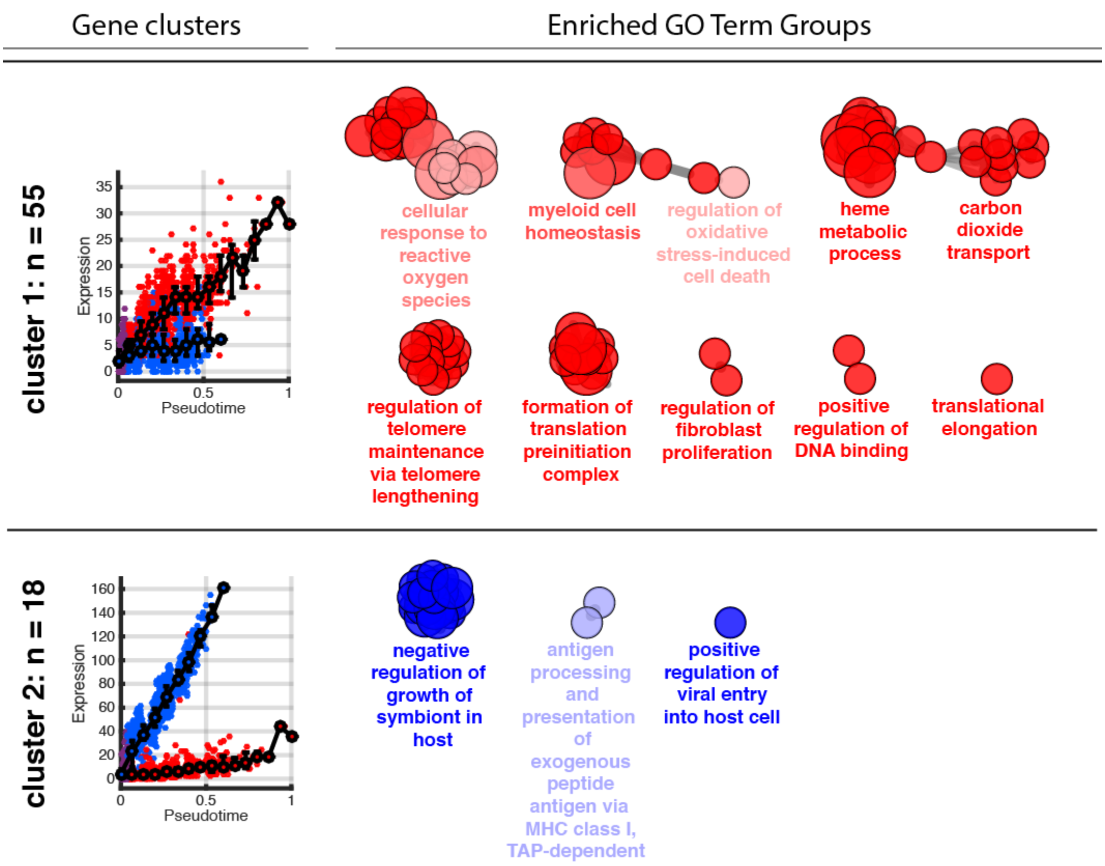


**Supplementary Figure 7.** GO enrichment analysis of branch-specific genes associated with the differentiation of Hematopoietic stem cells[29]. **LEFT**: The profile of representative genes in each cluster, where RED represents the cells assigned to Branch 1 and BLUE represents the cells assigned to Branch 2. **RIGHT**: Enriched GO term in each cluster, where each node represents a GO term related to the corresponding gene cluster, and all terms are functionally grouped by ClueGO**.** The most significant term is displayed below each group. Less cluster-specific terms are more transparent. Node size is proportional to significance (p-value). Only significant ($p > 0.05$) nodes are shown.

#### 3.7 Figures for demonstrating *Topographer's* robustness against the noise in a dataset

Although cell trajectories implied in a given set of data would be very complex and in spite of both cellular heterogeneity and gene expression noise in the dataset, *Topographer* can identify the backbones of de novo developmental trajectories from single-cell transcriptome data. **Supplementary Fig. 8-10** demonstrate the procedure of how *Topographer* identifies the backbone of cell trajectories from an artificial, noisy set of data, showing *Topographer*'s robustness ability in such identification.

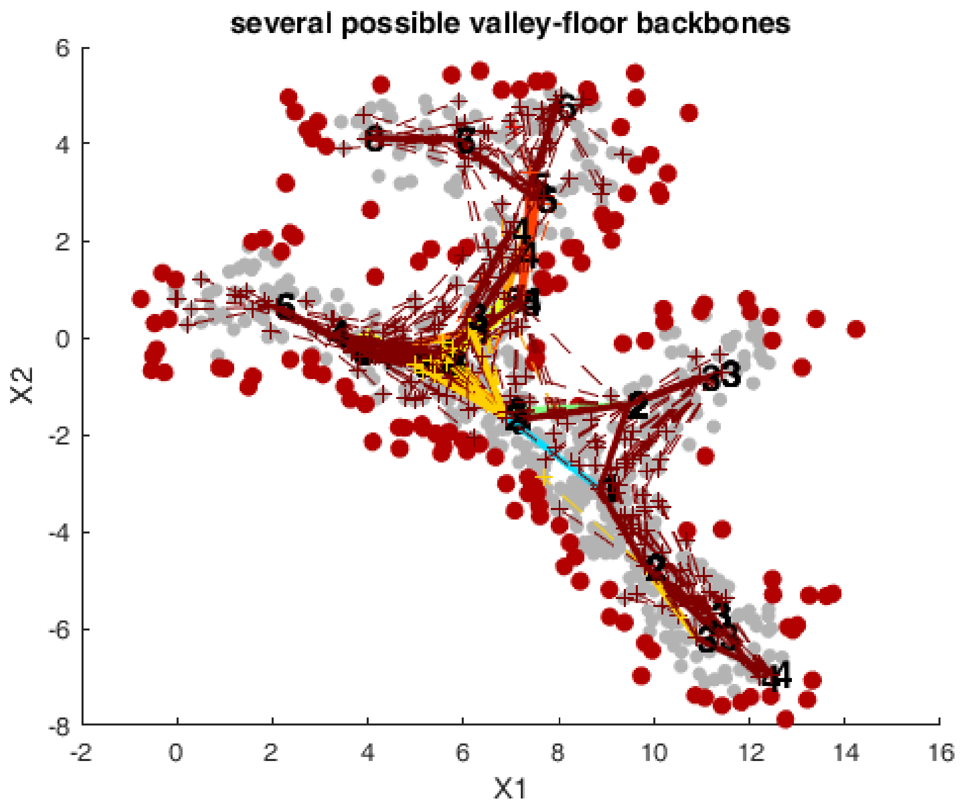


**Supplementary Figure 8** It is shown that *Topographer* finds many possible valley-floor backbones from a noisy set of data by repeating a clustering process for many times, where $x_1$ and $x_2$ represent two components.

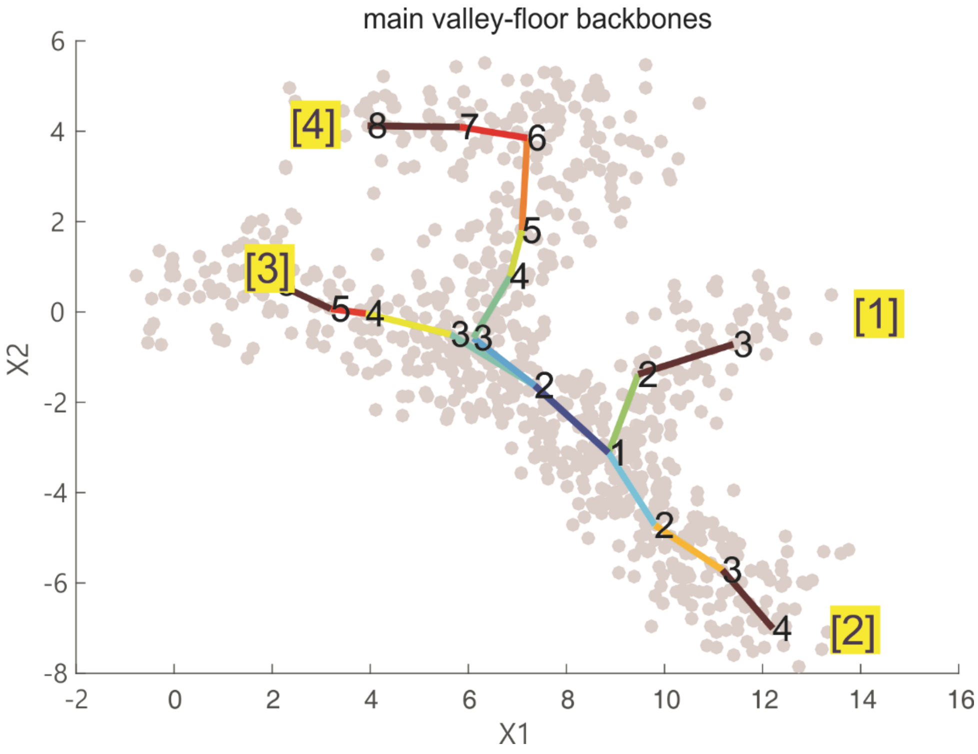


**Supplementary Figure 9** Four main valley-floor backbones identified by *Topographer*, where symbols [1] represents one main valley-floor backbone beginning at 1 and ending at 3, and symbols [2] represents another main valley-floor backbone beginning at 1 and ending at 4. Similar interpretations are for other two symbols: [3] and [4].

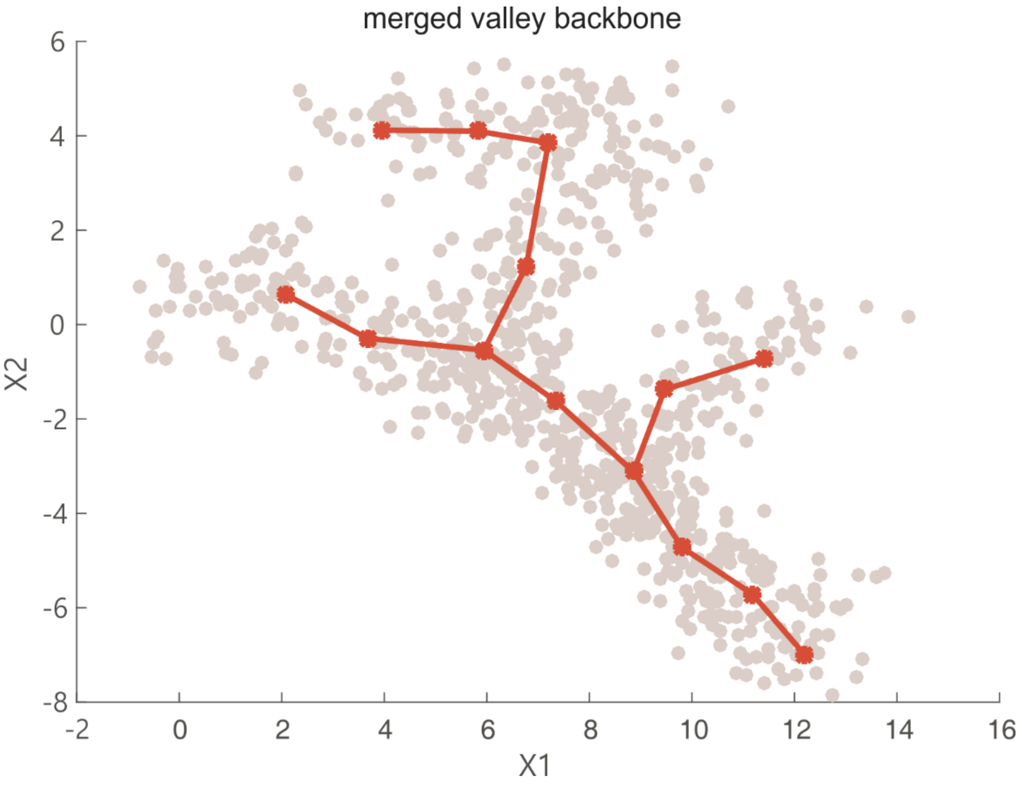


**Supplementary Figure 10** The final backbone obtained by merging the main valley-floor backbones in **Supplementary Fig. 9**.

#### 4. Analysis of a toy gene model

The following is a toy model for a differentiation regulatory network consisting of three pairs of antagonistic genes[30], which is used to generate an artificial set of data

$$
\begin{aligned}
\frac{dG_1}{dt} &= \frac{\alpha K_-^{\ h_-}}{K_-^{\ h_-} + G_2^{h_-}} - \delta G_1 + \xi_1(t) \\
\frac{dG_2}{dt} &= \frac{\alpha K_-^{\ h_-}}{K_-^{\ h_-} + G_1^{h_-}} - \delta G_2 + \xi_2(t) \\
\frac{dG_3}{dt} &= \frac{\alpha G_1^{h_+}}{K_+^{\ h_+} + G_1^{h_+}} \frac{K_-^{\ h_-}}{K_-^{\ h_-} + G_4^{h_-}} - \delta G_3 + \xi_3(t) \\
\frac{dG_4}{dt} &= \frac{\alpha G_1^{h_+}}{K_+^{\ h_+} + G_1^{h_+}} \frac{K_-^{\ h_-}}{K_-^{\ h_-} + G_3^{h_-}} - \delta G_4 + \xi_4(t) \\
\frac{dG_5}{dt} &= \frac{\alpha G_2^{h_+}}{K_+^{\ h_+} + G_2^{h_+}} \frac{K_-^{\ h_-}}{K^{h} + G_6^{h_-}} - \delta G_5 + \xi_5(t) \\
\frac{dG_6}{dt} &= \frac{\alpha G_1^{h}}{K_+^{\ h_+} + G_2^{h_+}} \frac{K_-^{\ h_-}}{K_-^{\ h_-} + G_5^{h_-}} - \delta G_6 + \xi_6(t)
\end{aligned}
\tag{24}
$$

where every $\xi_i(t)$ is the Gaussian white noise satisfying $\xi_i(t)\xi_j(t') = D\delta_{ij}\delta(t-t'), i = 1,\cdots,6$. Some parameter values are set as $\alpha = 250, \delta = 0.25, K_+ = 400, K_- = 200, h_+ = 20, h_- = 2$, $D = 100$. We simulated this set of stochastic differentiation equations using the Euler-Maruyama method[31], and the obtained results were shown in **Supplementary Fig. 11** and **12**.

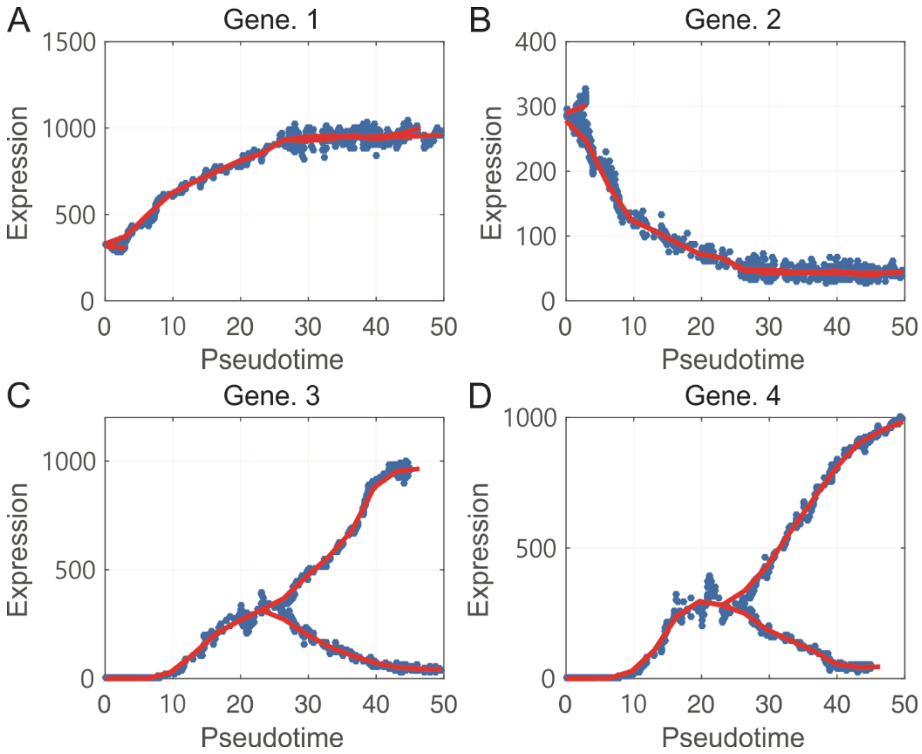


**Supplementary Figure 11** Shown are projected trajectories that *Topographer* constructed using an artificial set of data generated by a toy model (see **Eq. (24)**).

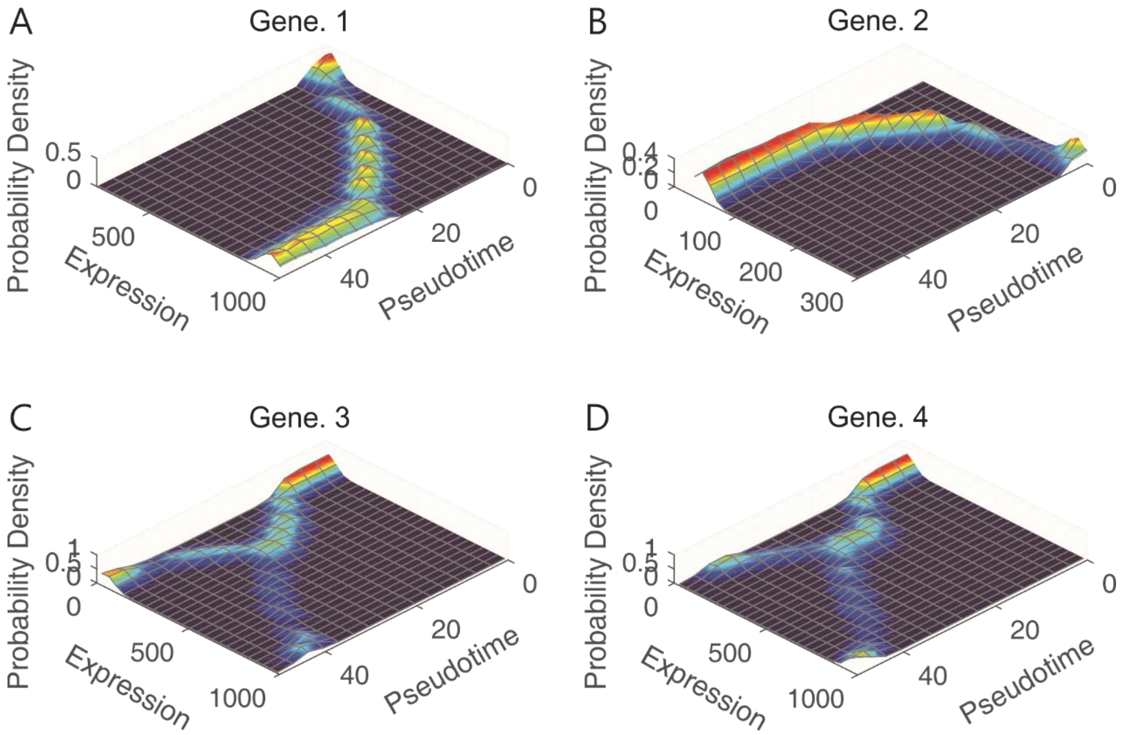


**Supplementary Figure 12.** Shown are three-dimensional trajectories that *Topographer* constructed using an artificial set of data generated by a toy model (also see **Eq. (24)**).

#### 5. Analysis of real data examples

Here, we used *Topographer* to analyze other two biological examples analyzed previously: the development of somatic stem cells[8] and the differentiation of Hematopoietic stem cells[29]. The aim is to further show the power of *Topographer*.

##### *5.1 The development of somatic stem cells*

Somatic stem cells contribute to tissue ontogenesis, homeostasis, and regeneration through sequential processes. Systematic molecular analysis of stem cell behavior is challenging because traditional approaches cannot resolve cellular heterogeneity or capture developmental dynamics. However, *Topographer* can provide a comprehensive resource of single-cell transcriptomes of adult hippocampal quiescent neural stem cells and their immediate progenitors.

In fact, by applying *Topographer* to a set of data on the development of somatic stem cells[8] (although the corresponding dataset contains only 132 cells and 23307 genes), we not only constructed an ‘intuitive’ (i.e., every cell is loaded with spatiotemporal information) developmental landscape (referring to **Supplementary Fig. 13A**) but also revealed stochastic dynamics of cell types by estimating both fate probabilities of cell types (4 cell types were found) and transition probabilities among them (referring to **Supplementary Fig. 13B**). Here we point out that these 4 found cell types have not been defined yet in the literature, so no names were indicated in the figure. **Supplementary Fig. 13C** showed evolutions of the transcription levels of four marker genes (Gfap, Apoe, Sox11 and Aldoc) along the identified cell trajectories.

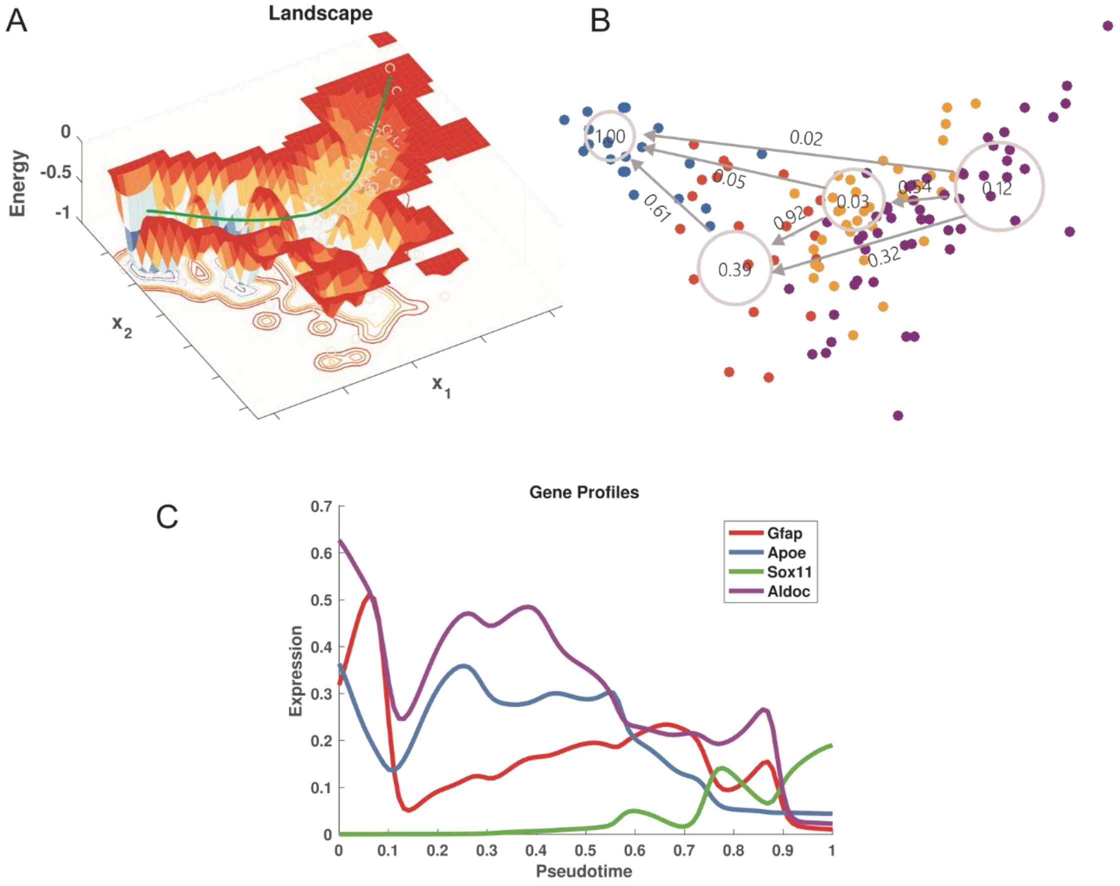


**Supplementary Figure 13.** *Topographer* constructs Waddington's potential landscapes and reveals stochastic dynamics of cell types, using a set of data on the development of somatic stem cells.

**(A)** A Waddington's potential landscape constructed by *Topographer*, where a thick green line represents the backbone of cell trajectories, and each small, grey circle represents one cell. $x_1$ and $x_2$ represent two components. The normalized potential is shown with depth of color representing the size of potential.

**(B)** Stochastic dynamics of cell types along branching trajectories, where both the fate probabilities of cell types (see circles with numbers) and transition probabilities among them are indicated.

**(C)** Evolutions of the expression levels of four marker genes (indicated by different colors) along the pseudotime (i.e., along the identified cell trajectory. See a thick green in (A)).

5.2 *The differentiation of Hematopoietic stem cells*

We retrieved a high-quality mass-parallel single-cell RNA-seq data on bone marrow cell differentiation[29], which contains a total number of more than 2,700 cells. We verified the globally low expressions of canonical lineage-determining transcriptional factor (TFs) of hematopoietic progenitors (Pu.1, Cebp-α, Cebp-β, Cebp-ε, Irf8, Gata1, Klf1, Gfi1b) in the pre-filtered dataset, consisting with Paul et al. 's view that the sorted cellular population belonging to early myeloid progenitors.

We selected a subset of cell states as 'initial cells' by gating on low expressions of canonical lineage committing TFs. With this set of 'initial cells', *Topographer* was then applied on the whole set of cells states after filtering the low expression genes, and dimensionally reduced to 50. As such, we constructed a trajectory with 2 branches, covering 60.04% and 59.75% of the complete population of cells sorted. The results were shown in **Supplementary Fig. 14** and **15**.

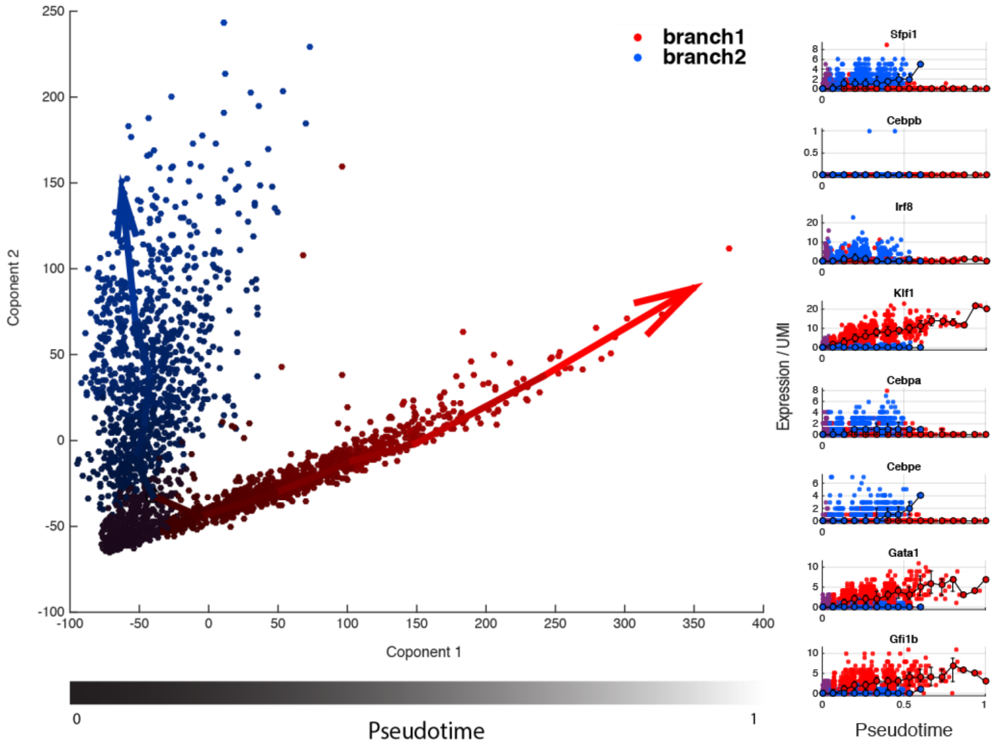


**Supplementary Figure 14. a)** Branching trajectories in the hematopoiesis dataset, identified by *Topographer* but projected on 2 dimensional PCA space, where RED represents the cells assigned to Branch 2 whereas BLUE represents the cells assigned to Branch 1, and the color mixed with gradient gray indicates the pseudotime. **b)** Profiles for a canonical lineage-determining transcriptional factor show globally low expressions, especially for Cebp-α and Irf8. Both branches show moderate (0 to 10 UMI) levels of some marker genes only in the later part of the branch. Dotted curves are calculated by sliding window average of the cells on each branch along the pseudotime. The same color represents the same cell type.

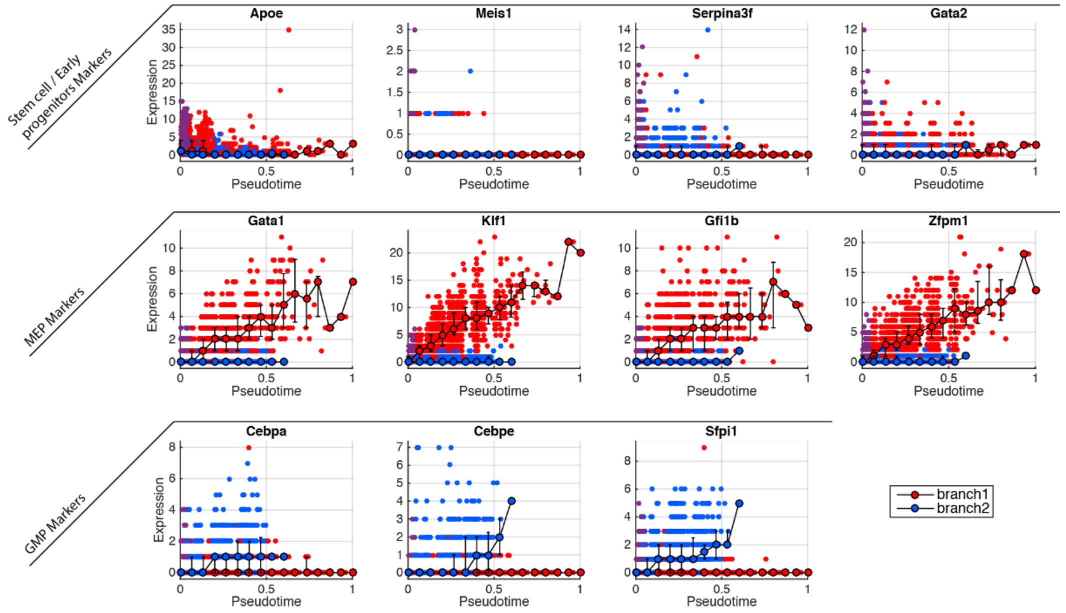


**Supplementary Figure 15.** Evolutions of selected marker genes along the pseudotime. First row: selected stem cells / early progenitor markers; Second row: selected MEP markers; Third row: selected GMP markers. Colors represent the same meanings as in **Supplementary Fig. 14**.
